# A pistil peptide toxin–pollen antidote system for reproductive barrier

**DOI:** 10.64898/2026.01.22.701211

**Authors:** Hiroki Miura, Kazuki Hirano, Takuya T. Nagae, Yoshinobu Kato, Taku Sasaki, Kenta Shirasawa, Shun Sakuraba, Tsukasa Matsuura, Sachiyo Soneda, Seiji Takayama, Sota Fujii

## Abstract

Prezygotic reproductive barriers in angiosperms are crucial for female-mediated selection of compatible male gametes. In this study, we identified a biological activity within the stylar tissues of *Arabidopsis thaliana* pistils that inhibits the elongation of heterospecific pollen tubes. Using genome-wide association studies, we identified a cysteine-rich peptide predominantly expressed in the style as the responsible barrier. This secreted peptide, named Femme fatale (FEM), inhibited pollen tube elongation of heterospecific species and some conspecific strains, both *in vivo* and *in vitro*. Importantly, FEM’s function was independent of the known reproductive barrier component SPRI1 that functions in the stigmatic papilla cell surface. This means two factors act spatially in sequence to create a multilayered reproductive barrier system against pollen in pistils. We also identified a secreted protein expressed in pollen, Homme capable (HOM), which is encoded by a gene next to *FEM*. HOM interacts with FEM to counteract its barrier effect and allow pollen tube elongation. The *FEM*–*HOM* locus has undergone frequent tandem duplications during Brassicaceae evolution. Dosage differences between the two factors can lead to reproductive incompatibility among lineages. Together, our findings suggest that FEM and HOM form a toxin-antidote system that helps establish reproductive barriers.

## Main

Prezygotic reproductive barriers are critical for speciation because they prevent the formation of undesirable hybrids. In angiosperms, open pistils are often exposed to interspecific pollen grains^1^. Studies show pollen transfer between different plant species is more common than expected^1,2^. Interspecific fertilization wastes resources that females would use for within-species reproduction. Thus, physiological barriers must form during pollen-pistil interactions. Reproductive interference between species is a key evolutionary force for these barriers. The study of prezygotic barriers has also become important in agroeconomics. Overcoming these barriers has been a long-standing interest, even discussed by Darwin in *The Origin of Species*.

Recent research on angiosperms has provided important insights into prezygotic pistil barriers. For example, the stigmatic *S*-locus receptor kinase (SRK) was long known for its role in the female determinant of self-incompatibility (SI) in Brassicaceae and was recently reported to contribute to the rejection of interspecific pollen grains^3^. In this system, Pollen coat protein B-class peptides induce rapid and selective hydration of conspecific pollen^3^. Both self- and interspecific incompatibility mechanisms are thought to recruit the receptor kinase Feronia (FER) to mediate reactive oxygen species signaling^3,4^. Rapid Alkalization Factor (RALF) peptides from pollen and stigma also interact with the FER receptor kinase complex to form a hybridization barrier^5^. These FER-dependent mechanisms show a connection between SI and interspecific barriers, which fits the classical SI x self-compatible (SC) unilateral incompatibility rule^3,6^. Such SI-related barriers also appear in Solanaceae. In these species, the pistil-side S-RNase and the pollen-side SLF complex components are directly involved in interspecific incompatibility^7,8^.

On the other hand, the pistil also has interspecific barrier systems unrelated to SI. One example in *Arabidopsis thaliana* is the stigmatic transmembrane protein Stigmatic Privacy 1 (SPRI1), which rejects heterospecific pollen from a broad range of species^9^. The SHI-family transcription factor SPRI2 regulates *SPRI1* and other genes needed for the stigmatic interspecific barrier^10^. More recently, the stigmatic cuticle layer of *Arabidopsis* was found to form a reproductive barrier^11^. In Solanaceae species, a farnesyl pyrophosphate synthase and a cysteine-rich protein SpDIR1L confer interspecific reproductive barriers^12,13^. In maize and wild teosinte relatives, silk- and pollen-expressed pectin methylesterases control reproductive barriers^14–16^. Other factors contribute to fertilization barriers in a Dobzhansky-Muller-like manner. These include the pistil-side pollen tube attractant LURE1 and its pollen-side receptor PRK6^17^. A subunit of the oligosaccharyltransferase complex, which helps inter-specific gametophyte recognition in *A. thaliana* and related species, may be another example^18^. Discoveries of these SI-independent mechanisms are beginning to reveal the multiplex nature of the pre-zygotic reproductive barrier in plants.

### Femme fatale forms a stylar reproductive barrier

During our quest to identify additional reproductive barrier factors, we observed that pistils of some *A. thaliana* accessions block elongation of *Erysimum allioni* pollen tubes*. E. allioni* is a relative of *A. thaliana* and belongs to the Brassicaceae family. This previously unrecognized incompatibility operates in the style, a pollen tube transmitting tract tissue within the pistil connecting the stigma to the ovary. For instance, six hours after pollination, pistils of *A. thaliana* Gel-1 or Col-0 accessions allowed robust *E. allioni* pollen tube elongation into the ovary zone. In contrast, most *E. allionii* pollen tubes were blocked at the styles of *A. thaliana* UduI1-11 or Me-0 accessions (Fig. 1a). We performed pollination assays at several time points to study the physiological nature of this barrier. When rejecting accession, Me-0 was the pistil-donor, *E. allionii* pollen tube elongation slowed at the stigma-style boundary (Fig. 1b,c). With non-rejecting accession Col-0 as the pistil-donor, *E. allionii* pollen tubes reached the ovary region with less slowdown (Fig. 1b,c). These results suggested that the style of some accessions, such as UduI1-11 or Me-0, suppresses heterospecific pollen tube elongation.

**Figure 1.**
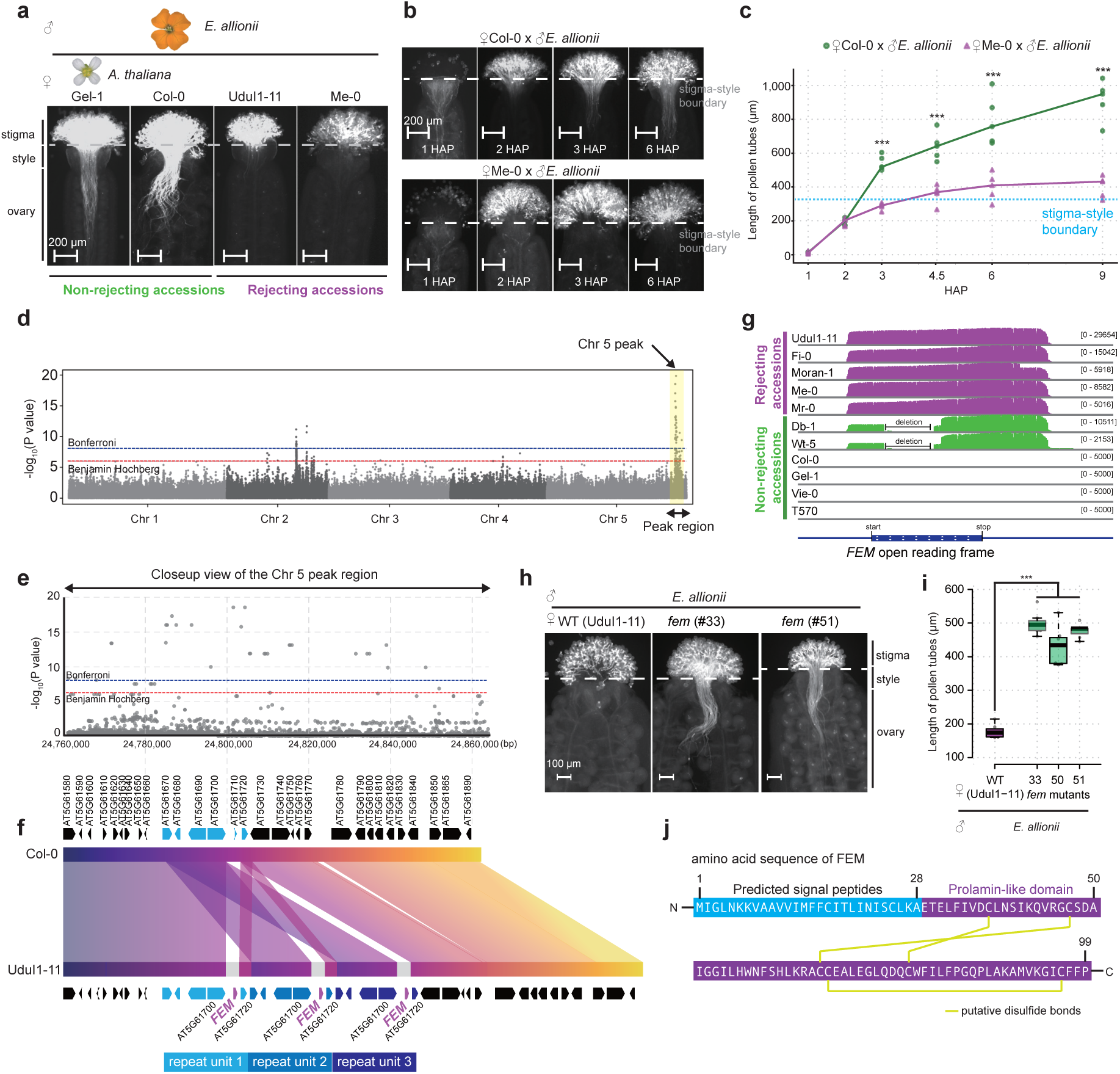
Identification of the *FEM* gene via GWAS. **(a**) Representative fluorescent images of aniline-blue-stained pistils fixed six hours after pollination with *E. allionii* pollen grains. Bars: 200 µm. (**b**) Representative fluorescent images of aniline-blue-stained pistils fixed at different hours after pollination (HAP) with *E. allionii* pollen grains. Bars: 200 µm. (**c**) Scatter plot of pollen tube length at different HAP. Lines pass through the mean at each HAP. *** indicate p < 0.005 by two-tailed Student’s *t*-test after Bonferroni corrections. (**d**) Manhattan plot of the GWAS. SNPs with minor allele frequencies > 0.15 are shown. The horizontal line indicates the nominal *P* < 0.05 threshold after Bonferroni (upper) and Benjamini-Hochberg (lower) corrections. (**e**) Close-up view of the Manhattan plot at the 24.76-24.86 Mbp region of chromosome 5. (**f**) Genomic synteny and the gene features in the *FEM* locus. Syntenic regions were detected by blastn. (**g**) IGV snapshot of *FEM* transcripts detected by the RNA-seq analysis. Unnormalized read counts are shown. (**h**) Representative fluorescent images of aniline-blue-stained pistils fixed six hours after pollination with *E. allionii* pollen grains. Bars: 100 µm. (**i**) Boxplot of pollen tube length. *** indicate p < 0.005 by two-tailed Student’s *t*-test after Bonferroni corrections. Center lines show the medians; box limits indicate the 25th and 75th percentiles; whiskers extend 1.5 times the interquartile range from the 25th and 75th percentiles; and data points are plotted as open circles. (**j**) Deduced amino acid sequence of FEM.

To find candidate loci governing the stylar reproductive barrier, we performed a genome-wide association study (GWAS) with 278 *A. thaliana* accessions (Supplementary Table 1). The pistils of the accessions were pollinated with *E. allionii* pollen, and then evaluated for their ability to prevent pollen tube entry into the styles. This identified a genetic locus on the long arm of chromosome 5 linked to the phenotype (Fig. 1d,e). However, the Col-0 transcriptome study found no strong candidate gene with high pistil mRNA expression^19^. We reasoned that this was because Col-0 lacks the reproductive barrier activity and the corresponding genetic factor (Fig. 1a). To explore a genetic factor specific to rejecting accessions, we used PacBio HiFi sequencing of UduI1-11 DNA to detect large genomic changes and novel genes. We found the GWAS peak subregion (about 24.76-24.86 Mbp on chromosome 5) was tandemly triplicated in UduI1-11 (Fig. 1f). Additionally, UduI1-11 did not have the AT5G61710 gene, but instead carried a gene absent from Col-0 within the triplicated region (Fig. 1f). We named this gene *Femme fatale* (*FEM*) for its role in preventing pollen tube elongation, as demonstrated below.

We next performed RNA-seq analysis using total RNA extracted from stylar tissues of 11 accessions that exhibited different *E. allionii* pollen tube rejection activities. We found that *FEM* in UduI1-11 was abundantly expressed in styles, and its mRNA expression was conserved among the accessions that exhibited elongation inhibition activities against *E. allionii* pollen tubes (Fig. 1g). Accessions lacking *E. allionii* pollen tube rejection activities either did not express *FEM* (e.g. Col-0) or carried genetic deletions within the open reading frame that most likely led to its loss of function (i.e. Db-1 and Wt-5) (Fig. 1g). To reveal its function in interspecific incompatibility, we generated *fem* mutants in the UduI1-11 background using the CRISPR/Cas9 system. For one of the *fem* mutant lines, we confirmed by whole-genome sequencing that all three *FEM* gene copies were knocked out, without deleting the proximal genes (Extended Data Fig. 1). The styles of the *fem* mutant lines permitted the entrance of *E. allionii* pollen tubes, while those of the wild-type UduI1-11 blocked them (Fig. 1h,i), indicating that FEM is responsible for the stylar pollen rejection activity. The *FEM* gene encodes a small cysteine-rich peptide that belongs to the group of Early Culture Abundant 1 (ECA1) gametogenesis-related family proteins, including *EGG CELL 1* (*EC1*) genes reported as required for fertilization^20^ (Fig. 1j and Extended Data Fig. 2a,b). In the phylogenetic tree of ECA1 proteins, FEM fell within a sub-clade with previously uncharacterized function (Extended Data Fig. 2a). Protein structure models predicted mature FEM to be composed of packed alpha-helical structures (Extended Data Fig. 2c). We here identified cysteine-rich peptide FEM as a factor that forms a stylar reproductive barrier.

### Multi-layered reproductive barriers

We then hypothesized that FEM, as a secreted cysteine-rich peptide, interacts directly with heterospecific pollen tubes and inhibits their elongation. To test this idea, we expressed and purified the secreted 6xHis-tagged FEM peptides (FEM-6xHis) in the *Pichia pastoris* system (Extended Data Fig. 2d,e,f). Elongation of *E. allionii* pollen tubes *in vitro* were significantly suppressed by treatment with purified FEM-6xHis in a dose-dependent manner (Fig. 2a,b). In contrast to *E. allionii,* elongation of UduI1-11 pollen tubes *in vitro* was unaffected by the treatment of FEM-6xHis, even at high concentrations (Fig. 2a,c).

**Figure 2.**
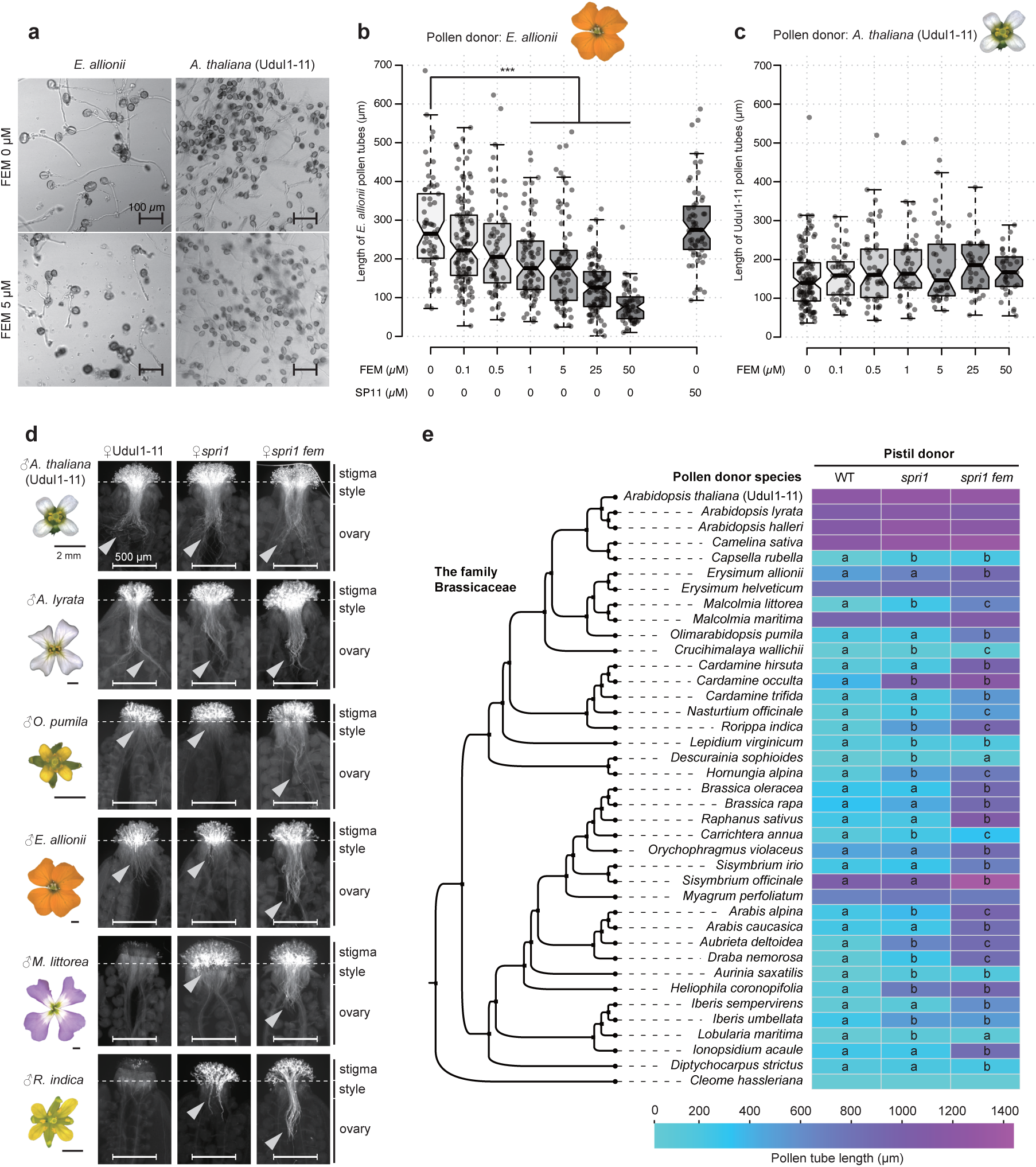
Functions of FEM. (**a**) Representative images of the *in vitro* pollen tube germination assays. (**b**) and (**c**) Boxplots of pollen tube length in *in vitro* pollen germination medium added with different concentrations of FEM. *** indicate p < 0.005 by two-tailed Student’s *t*-test after Bonferroni corrections. Center lines show the medians; box limits indicate the 25th and 75th percentiles; whiskers extend 1.5 times the interquartile range from the 25th and 75th percentiles; and data points are plotted as open circles. (**d**) Representative fluorescent images of aniline-blue-stained pistils fixed six hours after pollination. Bars: flower images, 1 mm; pistil images, 500 µm. (**e**) Cladogram of the Brassicaceae family adopted from ref^21^ and heatmap visualization of mean pollen tube length of each pollen pistil pair. Statistical groupings at the p < 0.05 significance level, determined by the Tukey Honestly Significant Difference (HSD) test, are indicated whenever a difference was found among the pistil donors.

Additionally, treatment of the SP11b peptide did not inhibit the elongation of *E. allionii* pollen tubes, indicating that the observed bioactivity is specific to the function of FEM (Fig. 2b). The above results suggest that FEM can directly inhibit the elongation of heterospecific pollen tubes.

We aimed to investigate the generality of the role of FEM and its relationship with SPRI1, the previously identified stigmatic reproductive barrier factor^9^. To explore this, we compared the abilities of the *spri1* single mutant as well as the *spri1 fem* double mutant generated in the UduI1-11 background, to reject pollen of 38 diverse species that belong to the Brassicaceae family (Extended Data Fig. 3). The SPRI1 barrier effectively blocked species such as *Malcolmia littorea* or *Rorippa indica*, since the *spri1* mutant permitted the pollen germination of these species when UduI1-11 rejected them at its stigma (Fig. 2d,e). However, pollen tubes of these species were blocked at the stigma-style boundary in the *spri1* mutant (Fig. 2d,e). Notably, pollen tubes of these species were able to penetrate to the ovary regions when the *spri1 fem* double mutant was used as the pistil donor (Fig. 2d,e).

This indicates that pollen tubes of *M. littorea* or *R. indica* are sensitive to both SPRI1 and FEM. On the contrary, pollen of species such as *Olimarabidopsis pumila* or *E. allionii* were insensitive to SPRI1 but were sensitive to FEM (Fig. 2d,e). The pollen of most species examined in this study was found to be rejected either by the function of SPRI1 or FEM, excluding those from close relatives of *A. thaliana* (e.g., *A. lyrata*, *A. halleri*, *Camelina sativa*), as well as pollen from an exceptionally distant species, *Myagrum perfoliatum* (Fig. 2e). Collectively, SPRI1 and FEM establish a multi-layered barrier that effectively responds to pollen grains originating from diverse evolutionary backgrounds.

### Pollen-side factor counteracts FEM

Upon challenging FEM functions against pollen of diverse origin, we unexpectedly observed that pollen tube elongation of some *A. thaliana* accessions was sensitive to FEM (Fig. 3a,b). Specifically, at four hours after pollination, pollen tubes of *A. thaliana* accessions such as Cdm-0 or Ge-0 were retained at the style in the presence of FEM (in the *spri1* mutant) but were able to reach the ovary region in the absence of FEM (in the *spri1 fem* mutant) (Fig. 3a,b). In contrast, pollen tubes of accessions such as Lis-3 or Qar-8a elongated regardless of the presence or absence of FEM (Fig. 3a,b). To eliminate the effect of SPRI1, we utilized the *spri1* mutant background for these experiments. We thus hypothesized that a pollen-side factor exists that counteracts the negative effect of FEM upon pollen tube elongation. We employed 414 *A. thaliana* natural accessions as pollen donors to assess their sensitivity to FEM by measuring pollen tube germination on pistils of both *spri1* and *spri1 fem* mutants of UduI1-11 (Supplementary Table 2). Using the relative pollen tube elongation ratio (*spri1 / spri1 fem*) as the input phenotype, we conducted GWAS to pinpoint the genetic locus associated with pollen tube sensitivity to FEM (Fig. 3c). A significant peak, collocated with the *FEM* region in the UduI-11 genome on chromosome 5, was identified from this analysis (Fig. 3c,d).

**Figure 3.**
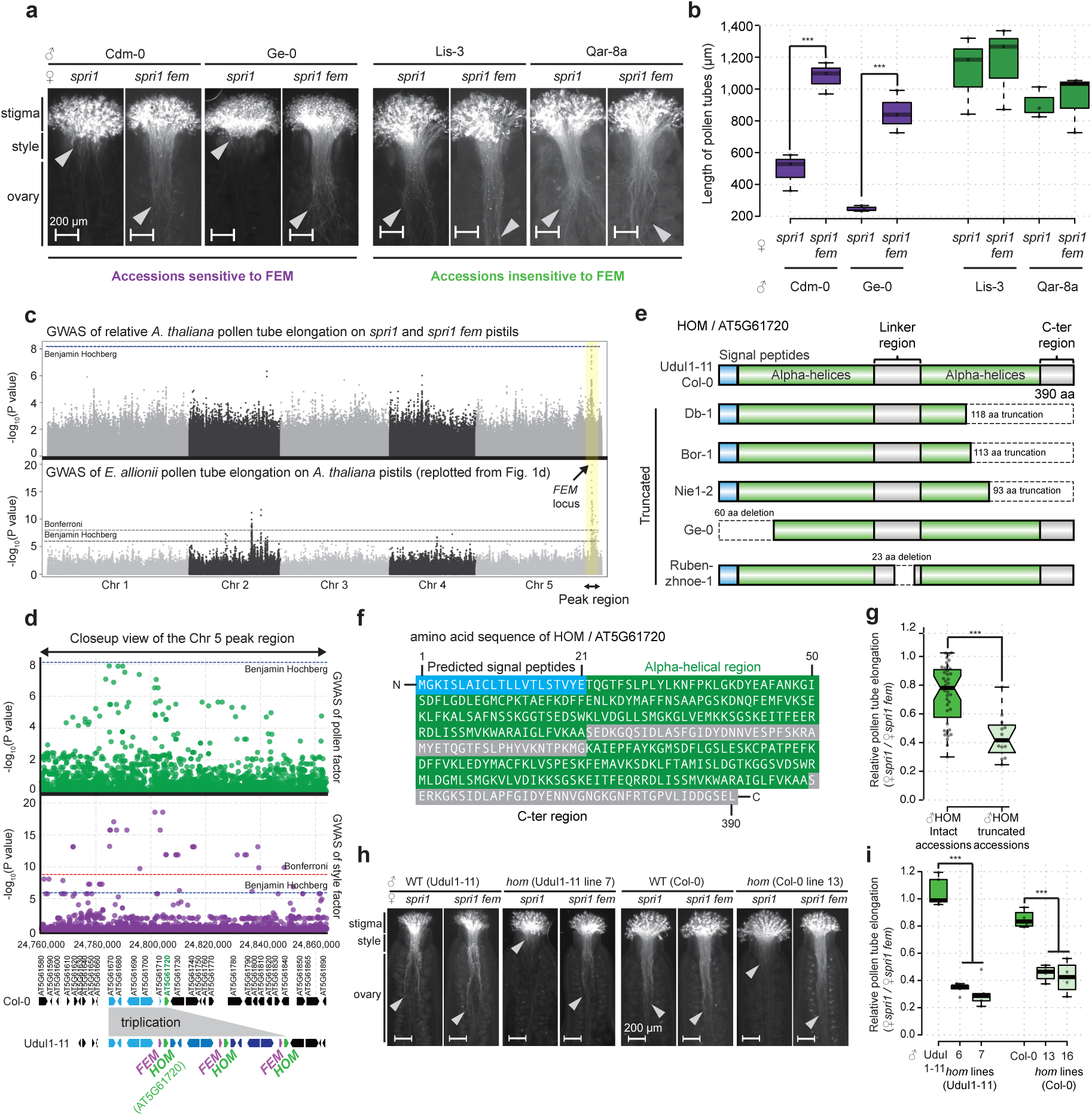
Identification of the *HOM* gene via GWAS. (**a**) Representative fluorescent images of aniline-blue-stained pistils fixed four hours after pollination. Bars: 200 µm. (**b**) Boxplot of pollen tube length. *** indicate p < 0.005 by two-tailed Student’s *t*-test after Bonferroni corrections. (**c**) Manhattan plot of the GWAS using the relative pollen tube penetration ratio (*spri1* / *spri1 fem*) of *A. thaliana* pollen as the input phenotype (upper). SNPs with minor allele frequencies > 0.15 are shown. The horizontal line indicates the nominal *P* < 0.05 threshold after Bonferroni (upper) and Benjamini-Hochberg (lower) corrections. GWAS of *E. allionii* pollen tube elongation on *A. thaliana* pistils was replotted from Fig. 1d for comparison (lower panel). (**d**) Close-up view of the Manhattan plot at the 24.76-24.86 Mbp region of chromosome 5. (**e**) Display of schematic protein secondary structure of HOM (AT5G61720) and natural variations causing possible truncations of the ORFs. (**f**) Deduced amino acid sequence of HOM (AT5G61720). (**g**) Boxplot of relative pollen tube elongation into *spri1* vs *spri1 fem* pistils. *** indicate p < 0.005 by two-tailed Student’s *t*-test. (**h**) Representative fluorescent images of aniline-blue-stained pistils fixed four hours after pollination. Bars: 200 µm. (**i**) Boxplot of relative pollen tube elongation into *spri1* vs *spri1 fem* pistils. *** indicate p < 0.005 by two-tailed Student’s *t*-test after Bonferroni corrections. For all box plots, center lines show the medians; box limits indicate the 25th and 75th percentiles; whiskers extend 1.5 times the interquartile range from the 25th and 75th percentiles; and data points are plotted as open circles.

We noticed that the gene adjacent to *FEM*, AT5G61720, was specifically expressed in mature pollen or anthers, as indicated by the *Arabidopsis* developmental transcriptome database^19^ (Extended Data Fig. 4a,b). AT5G61720 was predicted to encode a previously uncharacterized protein with Domain of Unknown Function 1216 (DUF1216). This protein was predicted to contain signal peptides at its N-terminus, and two clusters of alpha-helical regions (Fig. 3e,f, and Extended Data Fig. 4c,d). To discover natural deleterious mutations of this gene in *A. thaliana* accessions, we utilized the genomic sequence information of 156 *A. thaliana* accessions obtained by long-read sequencing technologies in past studies^22–24^, and in this study (Supplementary Table 3). We found that some *A. thaliana* accessions carry nonsense mutations within the AT5G61720 that may cause truncations to their open reading frames (Fig. 3e). In combination with the pollination data obtained for the GWAS, we found that pollen tube elongation of these accessions with truncated AT5G61720 were significantly more sensitive to FEM when compared to that of the accessions with intact AT5G61720 (Fig. 3g). We generated knockout mutants of this gene in Col-0 and UduI1-11 backgrounds (Extended Data Fig. 5) and pollinated the pistils of both *spri1* and *spri1 fem* mutants. As a result, the pollen tube elongation efficiency of the mutants of AT5G61720 was significantly reduced by the presence of FEM (Fig. 3h,i). This result indicated that pollen-expressed AT5G61720 attenuates the function of style FEM to persist pollen tube elongation. We named this pollen-specific gene *Homme capable* (*HOM*) for its bioactivity in counteracting the function of FEM in style. This series of evidence implied that the pistil peptide FEM and its tightly linked pollen factor HOM constitute a toxin-antidote-like system that contributes to the formation of a reproductive barrier.

### Molecular interaction of FEM and HOM

Both FEM and HOM were predicted to have N-terminal signal peptides, and thus, molecular interactions between the two factors were expected. We transiently expressed FEM fused with Venus at its N-terminus (Venus-FEM) and HOM fused with CFP at its C-terminus (HOM-CFP) in *Nicotiana benthamiana* tobacco leaves and observed under a confocal microscope (Fig. 4a and Extended Data Fig. 6a). First, Venus-FEM signals were detected in the region similar to AtPR3-mTurquoise2, the positive control for extracellular protein localization pattern (Fig. 4a). However, in contrast to AtPR3-mTurquoise2 signals that were evenly distributed to the extracellular regions, the Venus-FEM signals were found as non-continuous spotted patterns in the same region (Fig. 4a). When Venus-FEM and HOM-CFP were co-expressed, signals of both proteins co-localized in the extracellular regions, as the correlated spotted pattern (Fig. 4a and Extended Data Fig. 6b,c). To test the importance of FEM protein structures for the co-localization, we replaced two conserved cysteines of FEM with serine (C37S and C67S), and co-expressed the mutant Venus-FEMs with HOM-CFP. As a result, while signals of both C37S and C67S mutants accumulated as spotted patterns in the extracellular regions similar to the wild-type Venus-FEM, HOM-CFP signals were observed as dispersed in the extracellular region (Fig. 4a and Extended Data Fig. 6b,c). Therefore, the ability to co-localize with HOM was lost in the mutants of FEM. A luciferase complementation assay using split-Nanoluc moieties fused to HOM and FEM (Extended Data Fig. 6a) also showed that signals could be reconstituted by co-expression of the two fusion proteins (Fig. 4b,c). These observations suggested that both FEM and HOM are secreted into the extracellular space and may co-localize within the same molecular complex.

**Figure 4.**
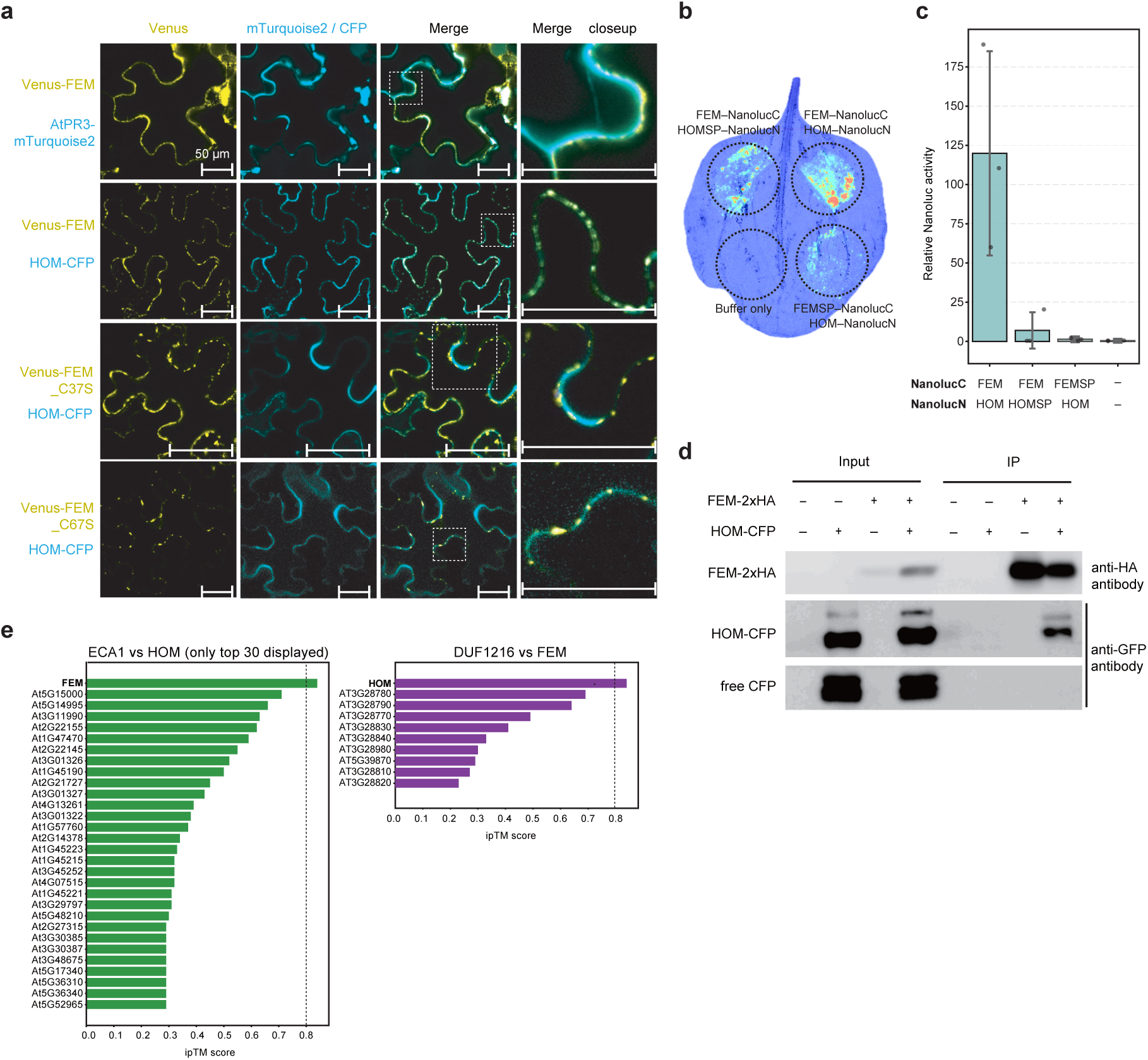
Molecular interaction of FEM and HOM. (**a**) Confocal fluorescence microscopy images of *N. benthamiana* leaf epidermal cells expressing designated pairs of proteins. Bars: 200 µm. (**b**) Representative images of the split Nanoluc assay using tobacco leaves. Bioluminescence was pseudocolorized by applying the lookup table “royal”. (**c**) Quantification of bioluminescence using the extracts of tobacco leaves. The relative activity of Nanoluc over firefly luciferase (driven by the *TGG1* promoter) ratio is displayed. (**d**) Co-immunoprecipitation analysis of FEM-2xHA and HOM-CFP. Input indicates total proteins extracted from Agrobacterium-infected *N. benthamiana* leaves. IP indicates fractions obtained by elution from the anti-HA agarose beads. (**e**) Bar graphs indicating ipTM scores obtained from the prediction of protein-protein interactions using Alphafold 3. Values greater than 0.8 are considered to represent confident, high-quality predictions^25^.

To further verify the molecular interaction of FEM and HOM, we co-expressed FEM fused with 2xHemagglutinin tag (FEM-2xHA) and HOM-CFP in tobacco leaves and conducted affinity purification using the anti-HA-agarose beads (Fig. 4d). The result showed that HOM-CFP, but not the putative free CFP probably produced by the cellular cleavage of HOM-CFP, was co-immunoprecipitated with FEM-2xHA (Fig. 4d). We also investigated whether such specific biomolecular interactions of FEM and HOM could be explained by their protein structural inferences. We evaluated the feasibility of various complexes formed by FEM and HOM homologues using AlphaFold protein structural prediction software^25^. The interface predicted template modelling scores (ipTM) for all combinations of 117 ECA1 proteins, including FEM, and 10 DUF1216 proteins, including HOM, were calculated based on the predicted complex structures. The ipTM scores reflect the “confidence” of the chain-chain interface predicted by AlphaFold3; in general, the value increases as the interface becomes more stable. The ipTM score of the protein complex structure predicted for FEM and HOM was 0.84, which ranked as the sixth among all 1,170 combinations tested, and they were each other’s highest ipTM-scoring partners (Fig. 4e, Extended Data Fig. 7). Above live-imaging, biochemical, and *in silico* studies suggested that FEM and HOM form a specific molecular complex, which further supports their role as the toxin-antidote pair.

### Evolution of the *FEM*–*HOM* locus

We were intrigued by the evolutionary dynamics of the FEM–HOM system, likely driven by the specific molecular interactions between the two factors. Tandem triplication of the *FEM*–*HOM* locus in UduI1-11 (Fig. 3d) prompted us to investigate the evolutionary trajectory of this genomic region. We therefore investigated the structural variation of the *FEM–HOM* locus across 156 *A. thaliana* accessions with whole-genome sequences available (Supplementary Table 3). Our analysis uncovered that among these, 27 accessions carried at least one copy of *FEM* and *HOM*, with both appearing functional (e.g., UduI1-11, Cvi-0) (Fig. 5a). Within these 27 accessions, 21 accessions carried multiple (two to five) copies of the tandemly duplicated *FEM–HOM* genetic locus (Fig. 5a,b). Additionally, 107 accessions carried *HOM* but lacked functional *FEM* (e.g., Col-0) (Fig. 5a). Furthermore, 22 accessions had lost both *FEM* and *HOM*, either through complete deletion of the gene locus or through apparent nonsense mutations (Fig. 5a). Notably, no accessions carried *FEM* without *HOM* (Fig. 5a). This was reasonable because the toxic function of FEM may lead to female self-sterility when HOM is absent.

**Figure 5.**
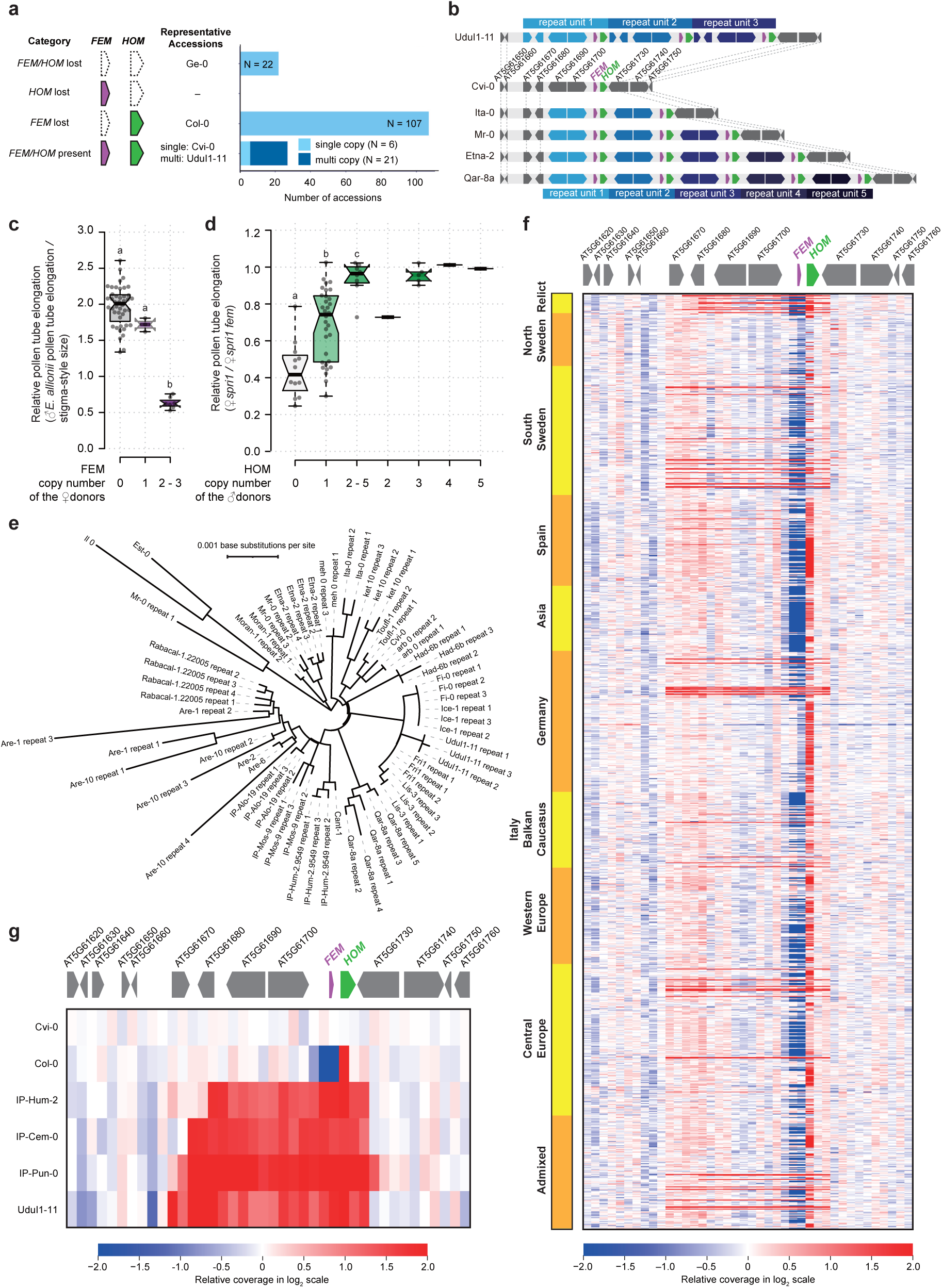
Molecular evolution of the FEM–HOM locus. (**a**) Summary of *FEM/HOM* carriers within the 156 accessions investigated. (**b**) Schematic display of the genomic structure and CNV of representative accessions carrying multiple copies of *FEM* and *HOM*. Single-copy genes flanking the *FEM*–*HOM* locus are connected to each other by broken lines. Boxplots of (**c**) relative pollen tube elongation of *E. allionii* into different *A. thaliana* accessions and (**d**) relative pollen tube elongation of different *A. thaliana* accession into *spri1* vs *spri1 fem* pistils. Statistical groupings at significance level p < 0.05 by Tukey HSD test are indicated. For (**d**), the statistical test has only been applied for categories with sufficient sample numbers (i.e., ‘0’, ‘1’, ‘2 - 5’). For both box plots, center lines show the medians; box limits indicate the 25th and 75th percentiles; whiskers extend 1.5 times the interquartile range from the 25th and 75th percentiles; and data points are plotted as open circles. (**e**) Evolutionary relationship of the 69 *FEM–HOM* nucleotide repeats inferred using the Neighbor-Joining method. (**f**) Heatmap of relative coverage depth of the 1,135 *A. thaliana* accessions to estimate the CNVs of the *FEM–HOM* locus. Accessions are in the order of ‘Admixture Group’ as inferred by ref^26^. (**g**) Display of the relative coverage heatmap for the selected accessions.

We sought to investigate the correlation between copy number variations (CNVs) in FEM–HOM and pollination phenotypes. Using the dataset we used for the GWAS (Fig. 1e), we first compared accessions that carry different copy numbers of *FEM* for their rejection activities against *E. allionii* pollen tubes in the style. As a result, accessions with two to three copies were active rejectors of *E. allionii* pollen tubes, while those with only one *FEM* copy showed little or no rejection activities against *E. allionii* pollen tubes (Fig. 5c). Similarly, we also revised our dataset, which we used for the GWAS to identify *HOM* (Fig. 3c), to investigate if *HOM* copy numbers affect pollen tube penetration efficiencies in the presence of FEM. We found that the genetic dosage of HOM is also positively correlated with its pollen tube competency, since accessions with multiple *HOM* copies elongated pollen tubes more efficiently into FEM pistils (Fig. 5d).

We next investigated the phylogenetic origin of the tandemly duplicated *FEM–HOM* locus. We constructed a phylogenetic tree from a sequence alignment of 69 “copies” of the *FEM-HOM* locus, extracted from 27 accessions that carried both *FEM* and *HOM* (Fig. 5a,b). We found that most of the time, nucleotide sequences were more conserved among copies than among accessions. For example, three copies found in UduI1-11 were phylogenetically closer to each other than the copies found in other *A. thaliana* accessions (Fig. 5e). This suggests that the UduI1-11 accession has experienced recent tandem duplication or gene conversion events of the *FEM–HOM* locus after its divergence from the other *A. thaliana* accessions. A similar situation is observed in copies of other accessions (Fig. 5e), suggesting that tandem duplications or gene conversion events may frequently occur at this genetic locus.

We further extended our investigation to the *Arabidopsis* 1001 genomes^26^. We expected that CNVs of the *FEM–HOM* locus of each strain could be estimated by mapping the short reads^26^ to the Cvi-0 genome, which carries a single copy of the *FEM–HOM* locus (Fig. 5a). From the coverage depth of the mapped reads, 60 *A. thaliana* accessions were estimated to carry multiple copies of the *FEM–HOM* locus, out of the 1,135 accessions investigated (Fig. 5f, Supplementary Table 4). Habitats of these accessions with multiple *FEM–HOM* copies were not limited to certain geographical groups inferred from the admixture analysis^26^ (Fig. 5f, Extended Data Fig. 8). The break points and lengths of the repeats were divergent among accessions, and at least four independent types of tandem duplications were found (Fig. 5g). These data suggested multiple parallel tandem duplications of the *FEM–HOM* locus have occurred during the evolution of *A. thaliana*.

Lastly, we performed comparative genomic analysis of the *FEM–HOM* locus in the Brassicaceae family using high-quality genome information (Extended Data Fig. 9). CNVs in species other than *A. thaliana* were also visible in the available genomic information. In *A. lyrata*, the MN47 accession carried two tandemly duplicated copies of *FEM–HOM*, whereas the NT1 accession carried five (Extended Data Fig. 9). Two accessions of *Camelia sativa* carried eight and ten copies of *FEM* and *HOM* (Extended Data Fig. 9). Therefore, tandem duplication was the conserved feature of the *FEM–HOM* locus in the family Brassicaceae.

## Discussion

In this study, we discovered the cysteine-rich peptide abundantly expressed in style of *A. thaliana* UduI-11 accession, FEM, as a factor that functions as reproductive barrier against *E. allionii* pollen (Fig. 1). FEM effectively suppresses elongation of pollen tubes from diverse Brassicaceae species, and together with SPRI1, it was shown to form a multi-layered interspecific reproductive barrier (Fig. 2). We also found that FEM can functionally operate as “toxin” even against intraspecific pollen, when its cognate “antidote” factor HOM is not functional (Fig. 3). We showed that *FEM*–*HOM* is a genetically linked toxin-antidote module that tandemly duplicated multiple times during the evolution of the Brassicaceae (Fig. 5).

There are many examples of cysteine-rich peptides involved in the arrest of plant cell growth. Examples of such regulations include the suppression of primary root elongation by the RALF peptide through interaction with FER^27^, and the inhibition of pollen tube elongation by tomato pollen SlPRALF^28^. In the meantime, competitive functions of pollen and pistil RALF peptides have been proposed in the regulation of pollen tube growth. Many pollen RALFs are found to promote pollen tube growth and elongation in both dicots and monocots^29,30^. Some pollen RALFs, such as RALF4/19, are known to act as ligands of CrRLK1L (*Catharanthus roseus* RLK1-like subfamily) receptors^31–33^, and/or retain pollen tube cell wall integrity via directly composing the extracellular polymers^34,35^. In some cases, antagonistic relationships between pollen and pistil RALFs are found.

Autocrine signaling maintenance of pollen cell wall integrity by RALF4/19 and its receptors BUPS1/2-ANX1/2 is competed by the female-expressed RALF34, and interference by this paracrine signal has been proposed to induce pollen tube burst^32^. It is possible that the FEM identified in this study could be interfering with pollen tube maintenance signaling pathways, thereby inhibiting cell elongation.

The molecular roles of HOM, on the other hand, remain rather speculative, as the biochemical function of the DUF1216 family has not yet been established. DUF1216 is a protein domain that is predominantly found in the proteomes of the Brassicaceae family^36,37^. They are one of the major proteins detected in the pollen grains by proteomic approaches, and it was predicted that this group of proteins may be involved in vesicle trafficking^38^. Typical DUF1216 proteins, including HOM, carry N-terminal signal peptides, with two patches of alpha-helical repeats connected by disordered linker protein sequences (Fig. 3e, Extended Data Fig. 4c). We detected molecular interactions in between FEM and HOM by imaging and biochemical approaches (Fig. 4), and this interaction may be direct as supported by the Alphafold 3 prediction (Fig. 4d, Extended Data Fig. 7). The flexible middle linker region of HOM may serve as the rheostat to enable specific physical interaction with FEM, although more mechanical studies are necessary to understand how this interaction evolved.

We propose that FEM–HOM is a tightly linked toxin-antidote module that cooperatively forms reproductive barriers. Such a relationship is reminiscent of the toxin-antidote system that triggers gamete-killing or developmental delays in rice^39,40^, maize^41^, fission yeast^42^, and nematode^43^. In these systems, a haplotype with selfish genetic elements, toxin and antidote functional units, can eliminate the haplotype lacking these elements, providing a simple explanation for the gene drive phenomenon^44^. In these evolutionary models, the acquisition of the antidote is proposed to precede the toxin, while its loss occurs after the toxin^44^. In line with these models, we observed that the majority of *A. thaliana* accessions, 134 out of 156 investigated, carried functional *HOM*, while only 27 accessions carried functional *FEM* (Fig. 5a). This suggested that HOM, the antidote, remained permissively after the loss of FEM toxin in many *A. thaliana* accessions. Loss of FEM–HOM most likely occurred as an evolutionary consequence of selfing in *A. thaliana.* The evolutionary origin of the *FEM*–*HOM* locus was considered to date back to the common ancestor of Brassicaceae, because the two genes are also linked in the genome of *Brassica rapa* (Extended Data Fig. 9), a species distantly related to *Arabidopsis* in this plant family. When tightly linked genetically, the quantities of these factors are maintained in balance, and discrepancies in their dosage (i.e., FEM > HOM) can result in incompatibility by suppressing pollen tube elongation in pistils (Fig. 5c,d).

The rapid evolutionary turnover of *FEM*–*HOM*, frequent genetic duplications and losses, and the opposing activities of male and female factors are characteristics that resemble other reproductive barriers found in plants. One example is the reproductive barrier in maize, controlled by the *Tcb1/Ga1/Ga2* genetic loci^14,45^. The *Ga1* locus carries different types of pectin methylesterases, the female factor ZmPME3 rejects incompatible pollen, and the male factor ZmGa1Ps-m confer pollen compatibility^14^. The S-RNase/SLF self-incompatibility system, shared by many angiosperm species, is another well-known example of reproductive barriers governed by a female toxin and a male antidote^46^. We here demonstrate the first example of a single locus-governed toxin-antidote-type interspecific reproductive barrier mediated by direct molecular interactions. It is possible that antagonism between pistil and pollen is a recurrent molecular evolutionary pattern in plants used for reproductive selection.

While interspecific pollination and reproductive interference are now being recognized as key evolutionary forces driving speciation, the importance of reproductive barriers is also being re-emphasized in this context^1^. Hybridization between different species can lead to inviable progeny due to post-zygotic incompatibilities^47^. Mechanisms to avoid loss of such resources may be important under these ecological and evolutionary selective forces, and recent studies have elucidated several important molecular factors that may serve as reproductive assurance of species^3,5,12,13,48^. Our study not only adds another repertoire to these factors but also discovered a previously unrecognized lock-and-key interaction between female and male, FEM and HOM. FEM and SPRI1 operate in different tissues within the pistil and together reject a wide range of heterospecific pollen (Fig. 2).

Moreover, it is likely that these factors also function independently of other factors, such as FER^3,5^, because SPRI1 has been shown not to be functionally related to SI^9^. Parallel requirements of these factors imply that reproductive assurance may be an important ecological issue for plants. The evolutionary selective pressure coming from floral pathogens that invade pistils^49^ could also be the biological significance of the female toxic factor, such as FEM. Alternatively, although less is understood about the evolutionary mechanisms of sexual selection in hermaphroditic plants, selection during pollen-pistil interactions may purge deleterious mutations by favoring males with higher genetic quality^50^. The FEM and HOM may play a central role in gamete selection by functioning as a toxin–antidote pair.

## Methods

### Plant materials

#### Plant materials and growth conditions

The GWAS seeds (Stock ID: N78942) were obtained from the *Arabidopsis* Biological Resource Center. Close relative species of *A. thaliana* were obtained as described in our previous study^9^. All plant materials were grown in mixed soil in a growth chamber under controlled conditions (light intensity of 120–150 µmol m^-2^ s^-1^; 14 h light/10 h dark cycles at 22 ± 2°C).

### Pollination experiments

Pollination experiments were performed as described previously^9^. Flowers were emasculated before anthesis. At anthesis, pistils were harvested, placed on 1% agar plates, pollinated, and incubated for the designated hours at 22 ± 2°C and 50 ± 5% humidity. In the pollinations, the entire stigmatic surface was covered with pollen grains. Pollinated pistils were fixed overnight at room temperature in ethanol/acetate (3:1, v/v), then heated to 60°C for 30 minutes in 1 M sodium hydroxide. The pistils were stained in 2% tripotassium phosphate/0.01% aniline blue for three hours at room temperature.

### Microscopic observation of pollen tubes and phenotyping

Pollen tubes on pistils stained with aniline blue were observed under an epifluorescent microscope as previously described^9^. For the GWAS using *E. allionii* as a pollen donor and *A. thaliana* natural variations as pistil donors, strains that inhibited *E. allionii* pollen tubes at their styles were regarded as “rejecting”, while those that allowed entrance of that into the ovary were regarded as “accepting”. For the GWAS using *spri1* and *spri1 fem* mutants (in UduI1-11 background) as pistil donors and *A. thaliana* natural variations as pollen donors, the length of pollen tubes was estimated from the aniline blue images. Fiji^51^ was used to measure pollen tube length in pistils.

### Genome-Wide Association Study

The GWAS was performed using the platform in the GWA-Portal website^52^. We used the 1001 imputed full-genome SNP set^26^ and the “accelerated mixed model”, both of which are available at the GWA-Portal. For the Manhattan plot, we removed SNPs with minor allele frequencies of < 0.15.

### Genome sequencing and assembly

We performed genome sequencing analysis of *A. thaliana* accessions Bil-5, Fi-0, Fri-1, Kni-1, Lis-3, Moran-1, Nie1-2, Ta-0, Tac-0, UduI1-11, Vie-0, Vimmerby and Wt-5 for use in the comparative genomic analysis. We also sequenced the *fem* mutant No. 33 (UduI1-11 background) to confirm the edited genome structure of the genetic line. Genomic DNA was extracted from the young leaves of these plants using Genomic Tip (Qiagen) and sheared to an average fragment size of 30 kb using Megaruptor 2 (Deagenode) in the Large Fragment Hydropore mode. The sheared DNA was used to prepare a HiFi SMRTbell library with the SMRTbell Express Template Prep Kit 3.0 (PacBio). The resultant library was separated on BluePippin (Sage Science) to remove short DNA fragments (<15 kb) and sequenced with SMRT cell 25 M on a Revio system (PacBio). The obtained HiFi reads were assembled with Hifiasm with the default parameters. Completeness of the assemblies was assessed using the embryophyta_odb10 data set with Benchmarking Universal Single-Copy Orthologs (BUSCO)^21^.

### RNA-seq analysis

RNA-seq was performed using Nanopore sequencing (Oxford Nanopore Technologies) as described previously^10^. Briefly, total RNA was extracted from the styles of 11 *A. thaliana* accessions (UduI1-11, Fi-0, Moran-1, Me-0, Mr-0, Db-1, Wt-5, Col-0, Gel-1, Vie-0, and T570) using the Maxwell Plant RNA Kit and the Maxwell RSC Instrument (Promega).

RNAseq libraries were constructed using the PCR-cDNA Barcoding kit (SQK-PCB109, Oxford Nanopore Technologies) following the manufacturer’s protocol. Sequence analysis was performed on the MinION instrument using the R9.4.1 version flow cell. The base calls were performed with Guppy (v5.0.11). Reads were mapped to the *A. thaliana* UduI1-11 genome using minimap2 (v2.24)^53^. Integrative Genome Viewer (IGV; v2.4.10)^54^ was used to visualize the mRNA accumulation.

### Genome editing

CRISPR/Cas9-mediated editing of *FEM, SPRI1,* and *HOM* was performed as described previously^10^ using the pHEE401 system, which expresses the *Cas9* gene specifically in the egg cell^55^. pHEE401E was a gift from Qi-Jun Chen (Addgene plasmid #71287). gRNA sequences were designed using the CRISPR-P program^56^. The gRNAs used for *FEM* were 5’-AACTCAATTAAGCAAGTCAG-3’, 5’-CGAAGCACTCGAAGGATTGC-3’; *SPRI1*, 5’-CGTATCTTCTATCTTGGCA-3’; and *HOM*, 5’-GCACTGTTTATGAAACACAA-3’, 5’-GCTTTCAACAGTAGTAAAGG-3’.

### Protein expression and purification using the *Pichia pastoris* system

A 255 bp gene fragment encoding the predicted mature sequence of FEM (encoding amino acid residues E29 to P99), a 2xGGGS linker, and a 6xHis tag, excluding the signal sequence predicted by SignalP^57^ and the stop codon, was inserted into the *Pichia pastoris* secretion expression vector pPIC9K (Thermo Fisher Scientific) at the NotI site. This plasmid was linearized by SalI digestion and introduced into *Pichia pastoris* GS115 by electroporation. Colonies carrying multiple copies of the expression cassette were selected on RDB medium and then inoculated onto YPD solid medium containing G-418 (Thermo Fisher Scientific) at 0.5 mg/mL. Cells were then cultured in 1.5 L of BMMY liquid medium under a rotary shaker at 22 °C for 6 days. Methanol at a final concentration of 0.5% was added every 24 hours to induce protein expression.

FEM with 2xGGGS and 6xHis added to its C-terminus (FEM-6xHis) was purified from the culture. The supernatant was obtained by centrifuging the culture medium at 3,500 × g, then filtering through Glass Microfiber Filters/Grade 13400 (Sartorius) and a Millipore Express PLUS Membrane Filter (Merck) using a Büchner funnel to remove the *P. pastoris* cells. This culture medium was concentrated to 150 mL (1/10 of the original volume) using the Vivaflow 50R (Sartorius) filtration unit, with a molecular weight cut-off of 5,000. It was then diluted 10-fold with binding buffer (20 mM KH₂PO₄, 13 mM K₂HPO₄, 20 mM imidazole, and 10 mM NaCl, pH 7.4) and was concentrated again twice. Ni-NTA Agarose (Qiagen) was added to this solution and incubated at 4°C with gentle agitation for 1 hour.

Ni-NTA Agarose beads were washed with the binding buffer three times. FEM-6xHis was then eluted from the Ni-NTA Agarose beads using elution buffer (20 mM potassium phosphate buffer (pH 7.4), 500 mM imidazole, and 10 mM NaCl). This solution was passed through a PD-10 desalting column (Cytiva) to remove salts and was buffered to 10 mM ammonium acetate. The solution was freeze-dried overnight to obtain purified FEM-6xHis.

### *In vitro* pollen germination assay

An *in vitro* pollen germination assay was performed as described previously^58^ with modifications. The pollen germination medium (PGM) contained 10% sucrose, 1 mM CaCl_2_, 1 mM Ca(NO_3_)_2_, 1 mM MgSO_4_, 0.01% boric acid, 1 mM PIPES (pH 7.0) in ultra-pure water. When using *A. thaliana* pollen, 10 μM epibrassinolide (epiBL; E1641, Sigma) was added to PGM to increase pollen germination efficiency^59^. About 15 flowers of *A. thaliana,* or 12 anthers of *E. allionii,* were collected into a 1.5 ml tube and vigorously vortexed in 500 μl PGM for 1 min to separate pollen grains from dehisced anthers. The tube was centrifuged at 5,000 × g for 1 min to pellet pollen grains. Collected pollen grains were resuspended in a fresh 200 μl PGM and transferred onto a glass-bottom dish (D11140H, Matsunami) for observation under the confocal laser scanning microscopy LSM880 (Carl Zeiss). The glass-bottom dishes were tightly sealed with parafilm after applying water droplets to their peripheries to prevent evaporation. Pollen tube germination was observed three hours after incubation at 22°C. Purified FEM-6xHis was dissolved in PGM and then added to the incubation medium. Chemically synthesized SP11b peptide was obtained as previously described^60^.

### Co-localization experiment design

The vector design for the co-localization experiment followed the previous study^61^. Primers used for PCR and their sequences are listed in Supplementary Table 5 and Supplementary Table 6, respectively. The Venus-FEM and HOM-CFP fragments generated by two-step fusion PCR were cloned into the pBI121 binary vector. To improve the protein expression efficiency, the 5’ UTR region of the *ADH* gene and the *AtHSP18.2* terminator sequences were added to the pBI121 binary vector, and Venus-FEM and HOM-CFP fragments were inserted in between these sequences. The sequence encoding the N-terminal 20 amino acids of AtPR3 (AT3G12500) fused to mTurquoise2, codon-optimized for expression in *N. benthamiana* (AtPR3-mTurquoise2), was synthesized by Genscript (Supplementary Table 6). The synthesized AtPR3-mTurquoise2 insert was cloned into the BsaI site of the pHREAC vector^62^ using the Golden Gate assembly kit (New England Biolabs). pHREAC was a gift from George Lomonossoff (Addgene plasmid #134908). The Quikchange mutagenesis protocol, as described previously^61^ was used to obtain binary vectors for the expression of Venus-FEM_C37S and Venus-FEM_C67S. The resultant plasmids were transformed into the *Agrobacterium tumefaciens* C58 strain.

### Split Nanoluc assay design

The vector design for the split Nanoluc assay followed the previous study^61^. Primers used for PCR and their sequences are listed in Supplementary Table 5 and Supplementary Table 6, respectively. The FEM-NanolucC (C-terminal 12 amino acids of Nanoluc fused to N-terminus of FEM) and HOM-NanolucN (N-terminal 159 amino acids of Nanoluc fused to HOM) fragments generated by PCR were cloned into the pBI121 binary vector. FEMSP-NanolucC (C-terminal 12 amino acids of Nanoluc fused to FEM signal peptide) and HOMSP- NanolucN (N-terminal 159 amino acids of Nanoluc fused to HOM signal peptide) were also generated as negative controls and cloned into the pBI121 binary vector. To improve the protein expression efficiency, the 5’ UTR region of the *ADH* gene and the *AtHSP18.2* terminator sequences were added to the 5’ and 3’ ends of these fragments, respectively. The resultant plasmids were transformed into the *Agrobacterium tumefaciens* C58 strain.

### Protein expression using the *N. benthamiana* system

An *Agrobacterium* infiltration experiment for the transient expression of proteins in *N. benthamiana* leaves was performed as previously described^61^. Briefly, each transformed *Agrobacterium* was cultured in the LB medium at 28°C for 16 h, pelleted and washed twice with 10 mM MES-KOH (pH 5.7) containing 10 mM MgCl_2_. Subsequently, the pelleted *Agrobacterium* was diluted to 10-fold the original culture volume with 10 mM MES-KOH (pH 5.7) containing 10 mM MgCl_2_ and 100 µM acetosyringone, and then infiltrated into *Nicotiana benthamiana* leaves. The infected leaves, 4 to 6 days post-infiltration, were used for fluorescent live-imaging or co-immunoprecipitation experiments.

### Fluorescent live-imaging

Tobacco epidermal cells were observed under the confocal laser scanning microscopy LSM880 (Carl Zeiss). The 440 nm Diode laser was used to detect the Fluorescence of mTurquoise2 and CFP. The 514 nm Argon laser was used to detect Venus signals. Images were taken using a Plan-Apochromat 40x/0.8 M27 immersion lens. Image acquisition was controlled using Zen 2.3 SP1 software (Carl Zeiss). Co-localization analysis of Venus–FEM and HOM–CFP was performed using the “coloc 2” plugin in Fiji^51^.

### Bioluminescence assay

Bioluminescence of *Agrobacterium*-infected tobacco leaves was measured using the Nano-Glo® Luciferase Assay System (Promega) according to the kit protocol. The substrate furimazine was diluted in the buffer provided with the kit and applied to the tobacco leaves. A *pTGG1*:*LUC* (Firefly luciferase) construct was co-expressed as an internal control for normalization. Leaves were observed with the chemiluminescence mode of ImageQuant™ LAS4000 (Cytiva). Bioluminescence images were merged onto bright field images by using the “Merge Channels” function in Fiji^51^. The merged image pseudocolorized by using the lookup table “royal”. To quantify bioluminescence, *Agrobacterium*-infected leaves were collected and pulverized in a mortar and pestle with substrate solution. The samples were centrifuged at 10,000 × g for 5 min to pellet leaf debris, and then the supernatant was collected to a 96-well plate. Bioluminescence was counted for 10 sec/well using a TriStar^2^ LB942 Modular Multimode Microplate Reader (Berthold).

### Co-immunoprecipitation analysis

Co-immunoprecipitation was performed similarly to our previous study^61^. *N. benthamiana* leaves were homogenized in homogenization buffer (50 mM HEPES-KOH (pH 7.4) containing 5 mM MgCl_2_, 2 mM MnCl_2_) with 20 mM sodium ascorbate and cOmplete EDTA-free protease inhibitor cocktail (Roche) added, using mortars and pestles. The debris was filtered using Miracloth (Merck). The filtered extract was centrifuged at 15,000 × g for 5 min, the supernatant was collected, and then centrifuged again at 17,000 × g for 30 min to obtain the soluble fraction. The soluble fraction was incubated with 5 µl of anti-HA beads included in the HA-tagged Protein purification kit (MBL Lifescience) on ice for 60 min.

The incubate was then transferred to the column included in the kit. The beads were washed three times with 200 µl of homogenization buffer containing 1% n-Dodecyl-β-D-maltoside. Twenty microliters of 2 mg/ml HA peptides was added to the beads to obtain the immunoprecipitated (IP) fraction. The eluted IP fraction was subjected to western blotting using anti-GFP-antibody (MBL Lifescience) or anti-HA-antibody (MBL Lifescience) as described in our previous study^61^.

### Retrieval of sequence information

Protein sequences of *A. thaliana* ECA1 and DUF1216 families were retrieved from the TAIR database. Protein sequences of FEM homologs in Brassicaceae were obtained from the Phytozome v13 database^63^. The *FEM–HOM* genomic regions of *A. thaliana* accessions were retrieved from sequences deposited by the pan-genome studies^22–24^ and also from the genome sequences obtained in this study, as explained above. The list of *A. thaliana* sequence accessions used in this study is provided in Supplementary Table 3. We investigated non-redundant 156 *A. thaliana* accessions. The blastn program^64^ was used to find the *FEM-HOM* regions in these sequences. The *FEM–HOM* syntenic regions of *A. lyrata* were obtained from chromosome-level assembly of two accessions^65^ and those of other Brassicaceae close relatives were obtained from the Phytozome v13 database^63^. The list of Brassicaceae close relatives used in this study and their sequence accessions is provided in Supplementary Table 7. Raw short read sequence reads of the *A. thaliana* 1,001 genomes deposited as PRJNA273563 bioproject were retrieved from NCBI SRA.

### Prediction of protein-protein interaction by Alphafold

The AlphaFold3 software^25^ was used to predict the structures of complexes. All possible combinations of the 117 ECA1 homolog proteins and 10 DUF1216 homolog proteins were considered, and their complex structures were predicted. We used the offline version of AlphaFold3 with default parameters. For each sequence pair, five independent structures were predicted, and the best structure with respect to the ranking score used in AlphaFold3 (determined from ipTM score, monomer prediction score, disorder, and clashes) was used for further analyses. The iPTM scores of these best complexes were used in the study. The PyMOL Molecular Graphics System, Version 3.0 (Schrödinger, LLC.), was used to visualize the predicted structures.

### Genome synteny analysis

Genomic synteny of the *FEM–HOM* regions within *A. thaliana* was searched by the blastn program^64^. Genomic synteny of the *FEM–HOM* regions among different species was searched by the tblastx program^64^. *Ab initio* gene predictions were performed by the Helixer v0.3.4 program^66^. Synteny plots were generated using custom Python scripts and the matplotlib library^67^, based on the blastn analysis output.

### Phylogenetic analysis

For the phylogenetic analysis of ECA1, putative signal peptides were predicted and removed by the SignalP v6.0 program^57^. Protein sequences were aligned using the MAFFT v7.490 program^68^, followed by manual correction of the alignment in the Jalview v2.11.4.1 program^69^. Sequences missing the conserved six cysteines or that appeared truncated were removed, resulting in an alignment of 120 proteins. Phylogenetic relationship was inferred by the Neighbor-Joining method using the MEGA v12 program^70^. The evolutionary distances were computed using the Dayhoff matrix. The partial deletion option was applied to eliminate all positions with less than 10% site coverage. Phylogenetic trees were visualized by the iTOL v7 server^71^.

For the phylogenetic analysis of *FEM–HOM* repeat regions, nucleotide sequences corresponding to AT5G61700 to HOM (AT5G61720) of the 69 copies were aligned using the MAFFT v7.490 program^68^, followed by manual correction of the alignment in the Jalview v2.11.4.1 program^69^. Phylogenetic relationship was inferred by the Neighbor-Joining method using the MEGA v12 program^70^. The complete deletion option was applied to eliminate positions containing gaps and missing data. Phylogenetic trees were visualized by the iTOL v7 server^71^.

### Analysis of CNVs

For studying copy number variations (CNVs) of the *FEM–HOM* locus, short read sequences of the 1,135 *A. thaliana* accessions were mapped to the genomic region from AT5G61620 to AT5G61760 of the Cvi-0 accession. The Cvi-0 accession was selected as the reference since it carries only a single copy of *FEM* and *HOM*. The minimap2 (v2.24) program^53^ was used to map short reads to the reference. CNVs of the *FEM–HOM* locus of the 1,135 *A. thaliana* accessions were estimated using the cnvkit (v0.9.11) program^72^. Coverage depth was calculated using the batch option. Relative coverage on a log_2_ scale was visualized as heatmaps using the seaborn library^73^ in Python.

### Statistical test and boxplot drawing

All statistical analysis was performed in R^74^. Boxplot drawing was done using BoxPlotR^75^.

## Data availability

RNA-seq data files were deposited at the National Center for Biotechnology Information Sequence Read Archive (NCBI-SRA) under BioProject ID PRJNA1343403. Raw sequence reads and the assembled sequences are available at DDBJ (BioProject accession number PRJDB38079) and Kazusa Genome Atlas (https://genome.kazusa.or.jp).

## Acknowledgements

We thank M. Okamura, T. Manabe, Y. Yamamoto, M. Nara, M. Ishii, K. Mori, M. Niidome, A. Yoshida, M. Saito for their technical assistance. This work was supported in part by a Grant-in-Aid for Transformative Research Areas (22H05172, 22H05174 to S.F. and S. Sak; 22H05172, 22H05181 to K.S.), Grants-in-Aid for Scientific Research (21H05030 to S.T.; 18H02456, 24K01692 to S.F.) by the Ministry of Education, Culture, Sports, Science and Technology of Japan (MEXT). This work was also supported by the PRESTO program (JPMJPR16Q8 to S.F.), the FOREST program (JPMJFR233S to S.F.) of the Japan Science and Technology Agency, the Suntory Rising Stars Encouragement Program in Life Sciences (to S.F.), and the Kazusa DNA Research Institute Foundation (to K.S.).

## Author Contributions

S.F. conceived the study. S.F., Y.K., T.S., and S.T. supervised the study. H.M. and S.F. wrote the manuscript with inputs from all other authors. H.M., K.H., T.T.N., S.Son., and S.F. performed pollination experiments. H.M., K.H., T.T.N., and S.F. conducted the GWAS. K.S. and T.M. conducted the genome sequencing analysis. H.M. and S.F. performed the RNAseq analysis. H.M., K.H., T.T.N., and S.Son. generated the transgenic lines. H.M., K.H., Y.K., and S.Son. conducted protein expression experiments using the *Pichia* system. Y.K., T.S. and S.Son. conducted protein expression experiments using the tobacco system. S.F. conducted the *in vitro* pollen germination assay. S.Son. and S.F. conducted the live-imaging experiment. T.S., S.Son. and S.F. conducted the split Nanoluc assay. Y.K. conducted the co-immunoprecipitation experiment. S.Sak. ran the Alphafold 3 prediction, collected the ipTM scores, and supervised the structural prediction studies. T.S. and S.F. conducted the comparative genomics analysis.

## Figure Legends of Extended Data

**Extended Data Figure 1. Genomic structure of the *FEM* locus of the *fem* mutant**

(**a**) Schematic model of the *FEM* locus genome structures of the *fem* mutant. A large part of repeat unit 2 was deleted in the *fem* mutant. Two copies of the *FEM* sequences remained in the *fem* mutant. (**b**) Genetic sequence of the two *FEM* copies of the *fem* mutant. Both copies were expected to have lost function, either via a genetic deletion or a one-base-pair insertion.

**Extended Data Figure 2. Phylogeny and protein profiles of FEM**

(**a**) Phylogenetic tree of the ECA1 family proteins inferred using the Neighbor-Joining method. The subfamily affiliation of cysteine-rich proteins is indicated as CRPXXXX. Robustness of the branches as inferred from the bootstrap analysis is indicated by circles. (**b**) Deduced amino acid sequence of FEM adopted from Fig. 1j. (**c**) Tertiary protein structure of FEM without signal peptides predicted by Alphafold 2. (**d**) Diagram of vector design of FEM-6xHis expression in *Pichia pastoris* system. (**e**) Scheme for affinity purification of FEM-6xHis. (**f**) Silver-staining of purified FEM-6xHis protein electrophoresed on an SDS-PAGE gel.

**Extended Data Figure 3. Pollination results of different Brassicaceae species**

Boxplots of pollen tube length of each species pollinated on WT (UduI1-11), *spri1,* and *spri1 fem*. Center lines show the medians; box limits indicate the 25th and 75th percentiles; whiskers extend 1.5 times the interquartile range from the 25th and 75th percentiles; and data points are plotted as circles. Statistical groupings at the p < 0.05 significance level, determined by the Tukey Honestly Significant Difference (HSD) test, are indicated whenever a difference was found among the pistil donors.

**Extended Data Figure 4. mRNA expression and protein structure profiles of HOM**

mRNA expression patterns of the *HOM* gene (AT5G61720) are displayed as (**a**) a visible diagram adopted from the TAIR database and (**b**) read counts extracted from the transcriptomic database^19^. (**c**) Deduced amino acid sequence of HOM (AT5G61720) adopted from Fig. 3f. (**d**) Tertiary protein structure of HOM without signal peptides predicted by Alphafold 3.

**Extended Data Figure 5. Genomic structure of the *FEM–HOM* locus of the *hom* mutants**

(**a**) Schematic model of the *FEM–HOM* locus genome structures of the *hom* mutants generated in both Col-0 and UduI1-11 backgrounds. The entire repeat unit 2 and a large part of repeat unit 2 were deleted in the *hom* mutant in the UduI1-11 background. Consequently, only one copy of the *HOM* sequence remained in this mutant. (**b**) Schematic structure of the *HOM* gene. (**c**) Genetic sequence of the *HOM* gene of the *hom* mutant generated in the Col-0 background. (**d**) Genetic sequence of the *HOM* gene of the *hom* mutant generated in the UduI1-11 background.

**Extended Data Figure 6. Co-localization analysis of Venus–FEM and HOM–CFP**

(**a**) Schematic diagram of the construct design of the co-localization, split Nanoluc, and Co-IP experiments. (**b**) Representative scatter plot outputs of co-localization analysis by the “coloc 2” plugin in Fiji. Scales are in arbitrary units (a.u). (**c**) Boxplot of Pearson’s R values calculated for all combinations. Experiments were repeated independently three times. Center lines show the medians; box limits indicate the 25th and 75th percentiles; whiskers extend 1.5 times the interquartile range from the 25th and 75th percentiles; and data points are plotted as circles.

**Extended Data Figure 7. Prediction of protein-protein interaction probabilities by Alphafold 3**

(**a**) Scheme for predicting protein-protein interaction probabilities of ECA1 and DUF1216 family proteins by Alphafold 3. (**b**) Heatmap of ipTM scores calculated for all 1,170 pairs. (**c**) Histogram showing frequency of ipTM scores calculated for all 1,170 pairs. (**d**) The best protein complex model for FEM (in magenta) and HOM (in green).

**Extended Data Figure 8. Geographical distribution of *A. thaliana* strains carrying multiple copies of *FEM–HOM* genes**

Distribution of 1,135 *A. thaliana* accessions and their relative coverage depth of the *FEM* genetic region (**a**) worldwide, (**b**) in Europe-Africa, and (**c**) in central Europe.

**Extended Data Figure 9. Gene alignment, structure, and synteny of the genomic region surrounding the *FEM*–*HOM* locus in different Brassicaceae species**

(**a**) Gene alignment synteny of the *FEM–HOM* locus in *A. thaliana*, *A. lyrata,* and *C. sativa*. (**b**) Gene alignment synteny of the *FEM–HOM* locus, including that of *B. rapa,* which spans a relatively broader region.

