## Extended Data Figures for "A pistil peptide toxin–pollen antidote system for reproductive barrier"

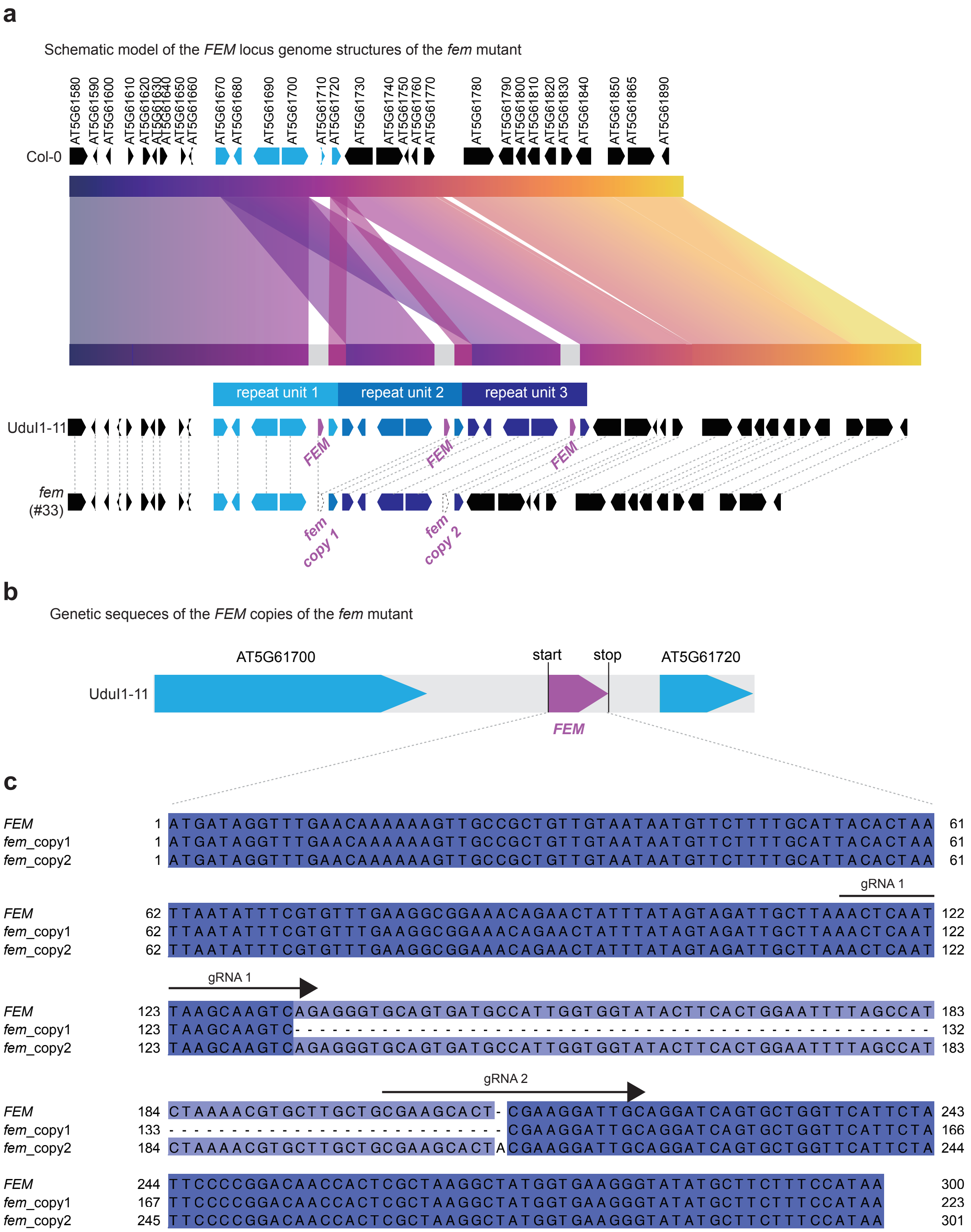


**Extended Data Figure 1**

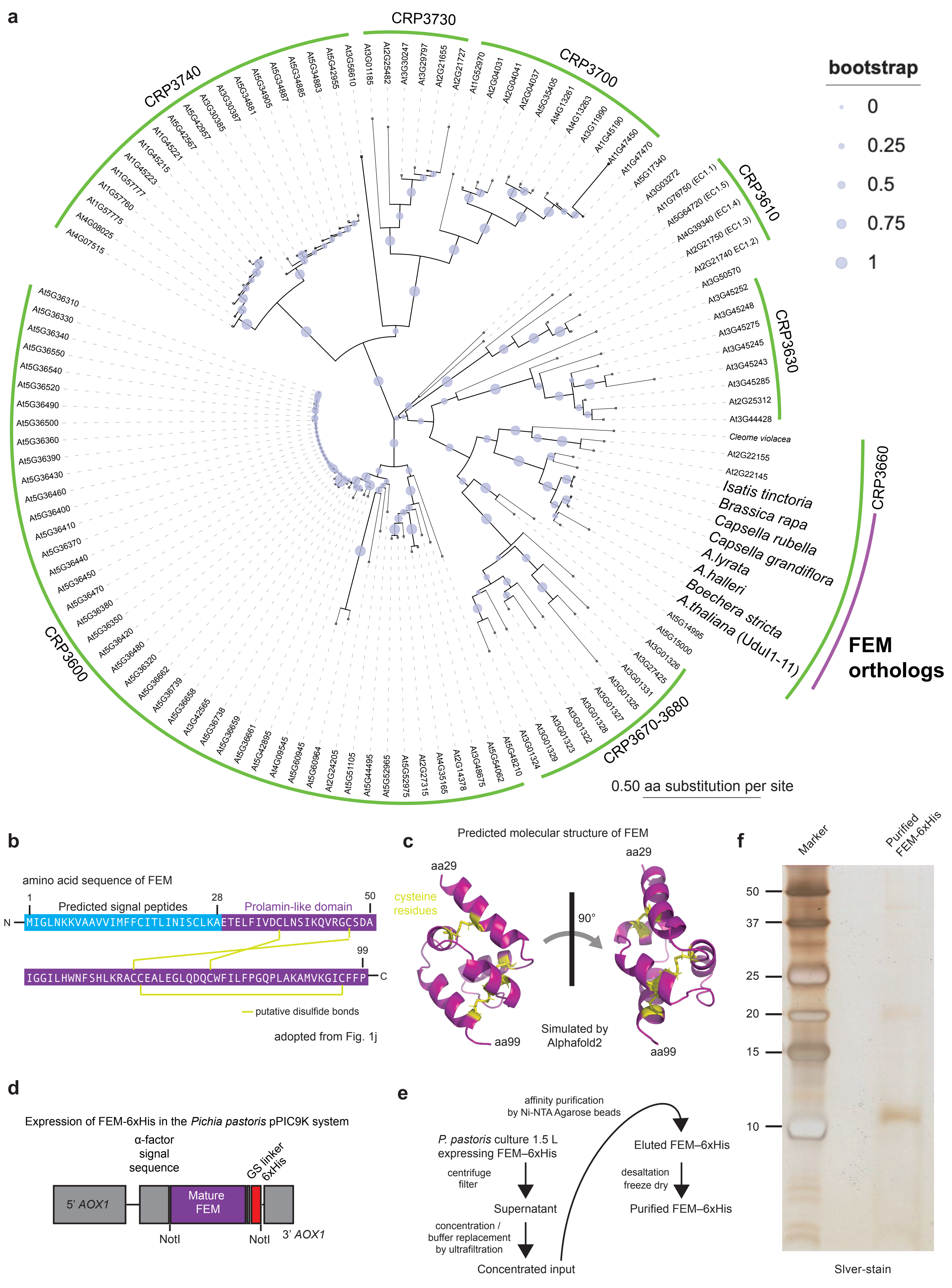


**Extended Data Figure 2**

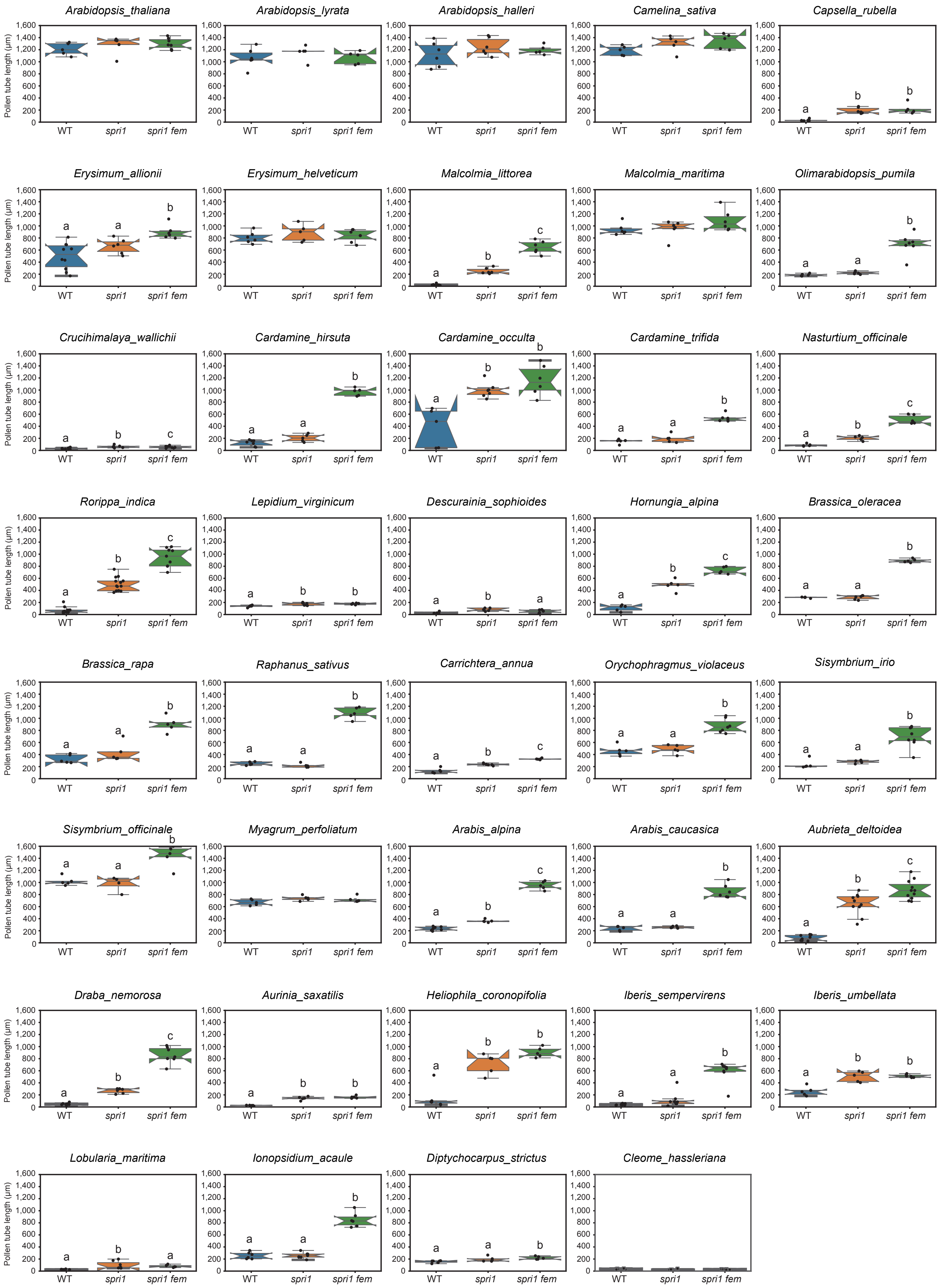


### Extended Data Figure 3

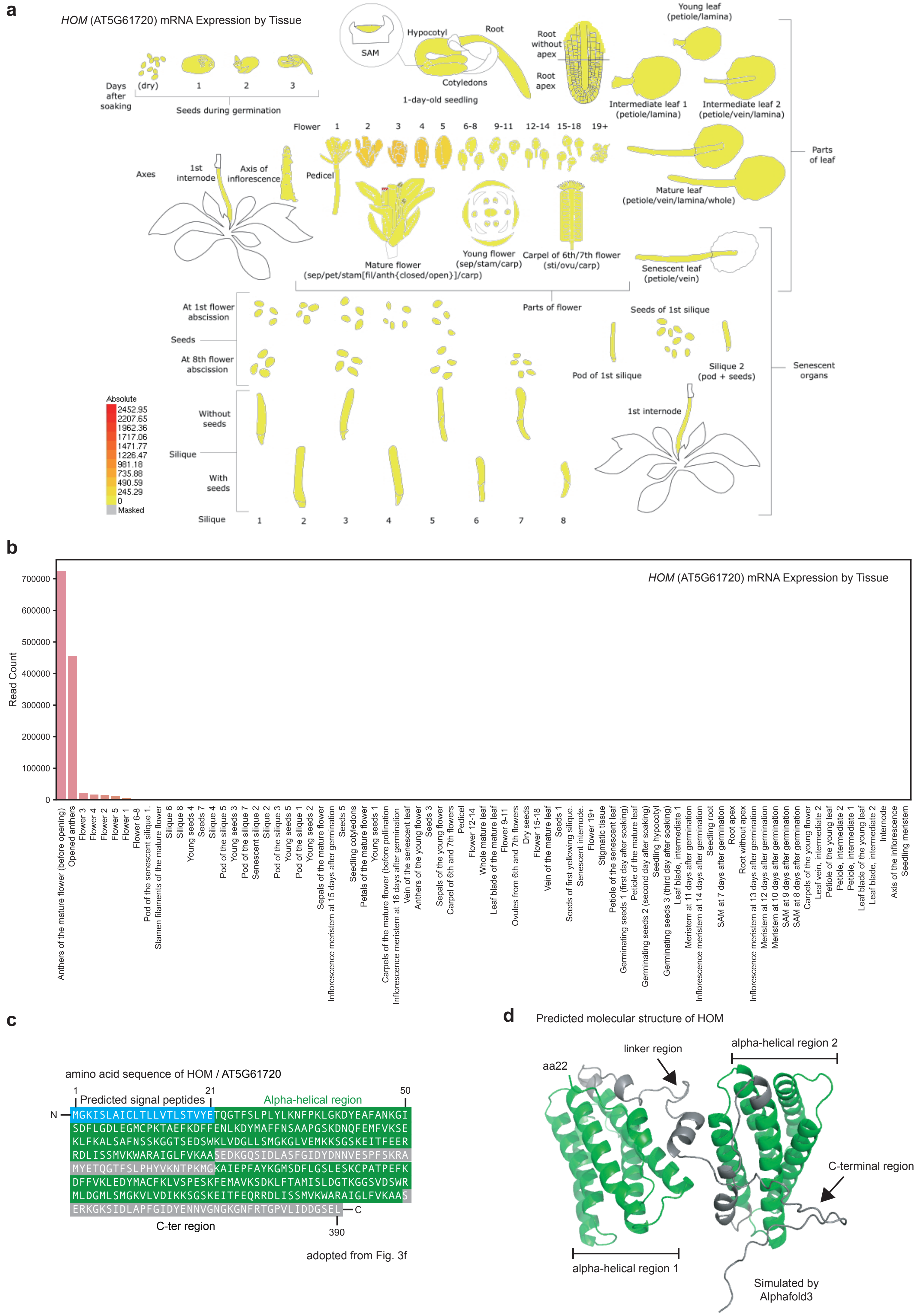


**Extended Data Figure 4**

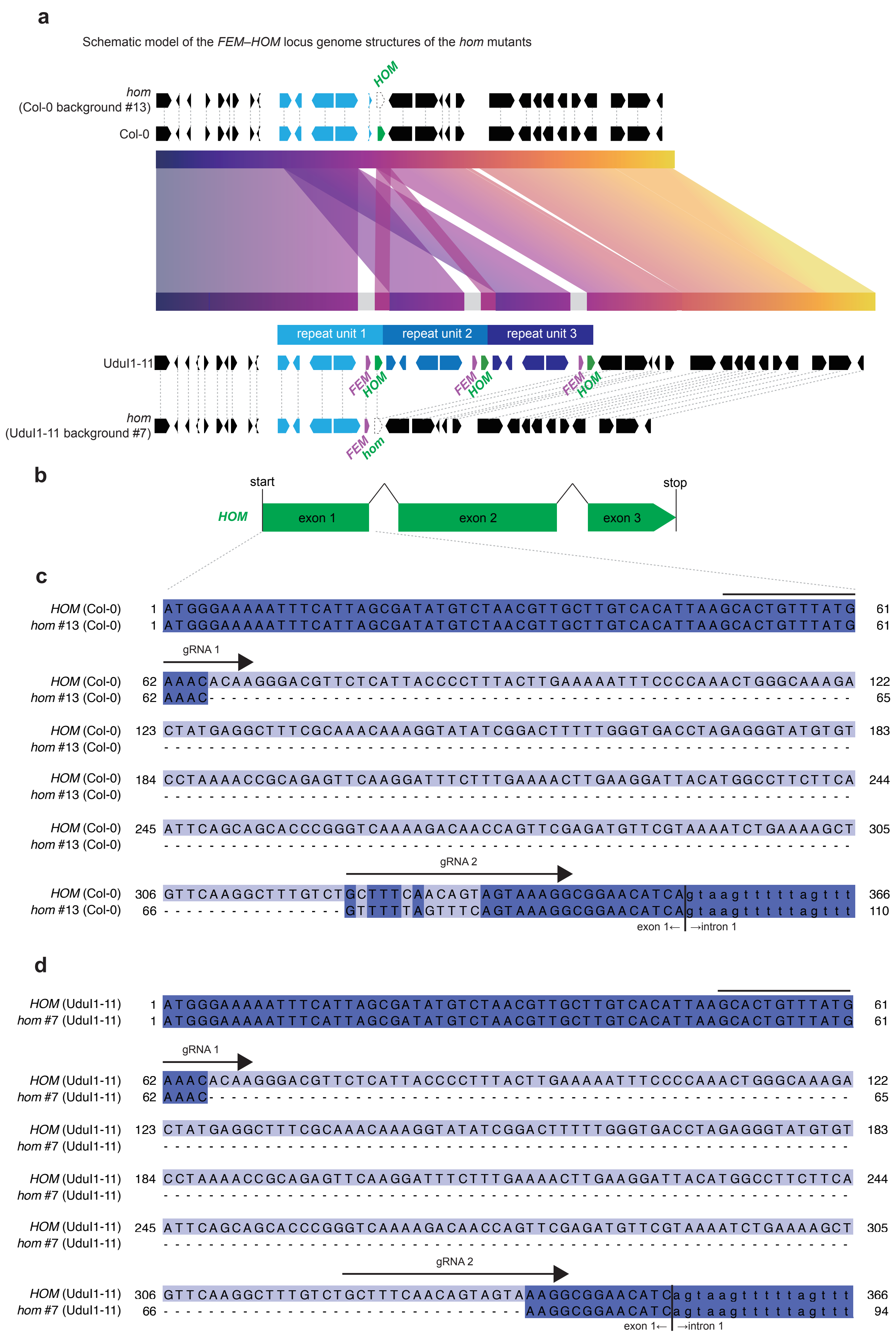


**Extended Data Figure 5**

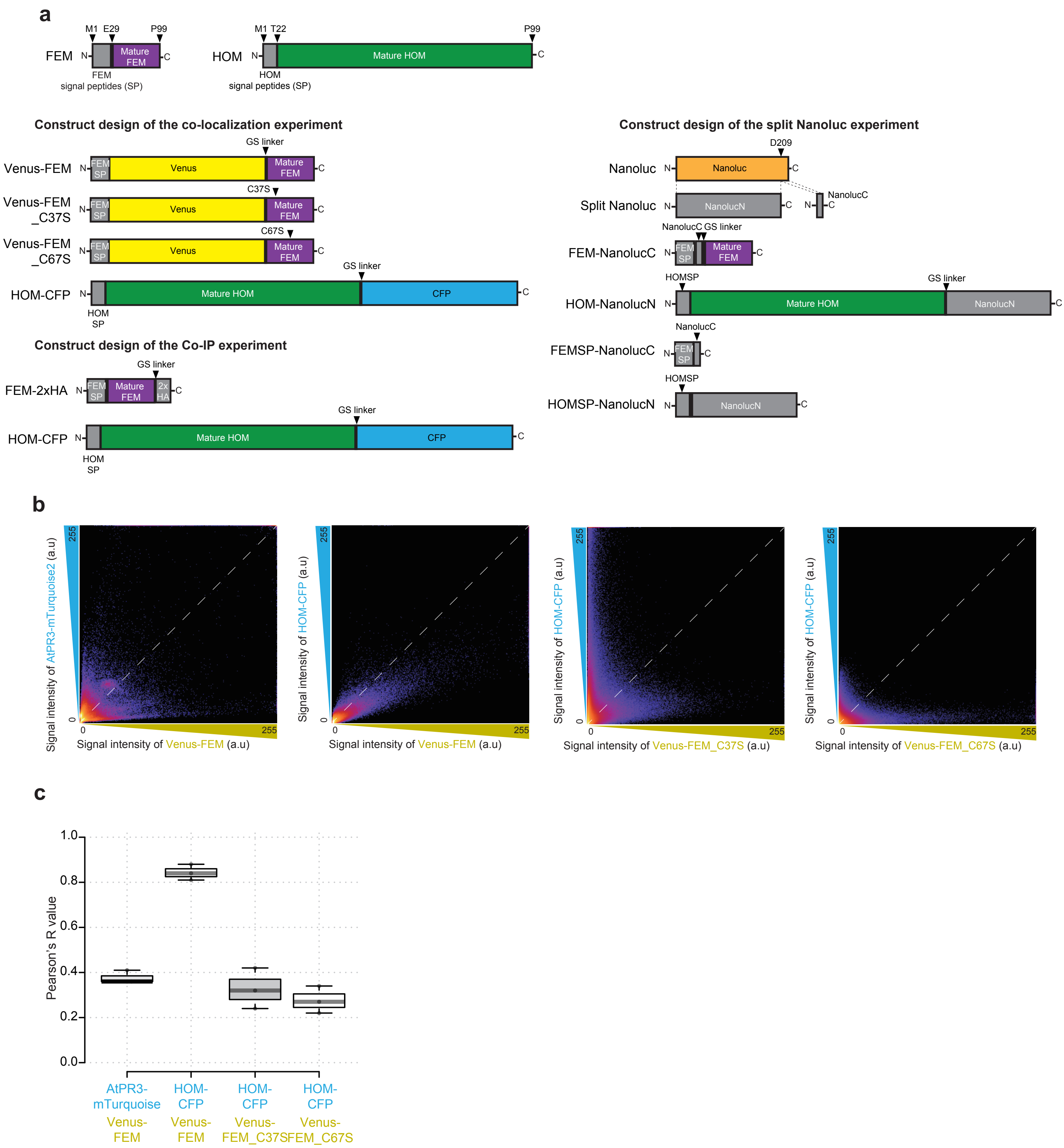


**Extended Data Figure 6**

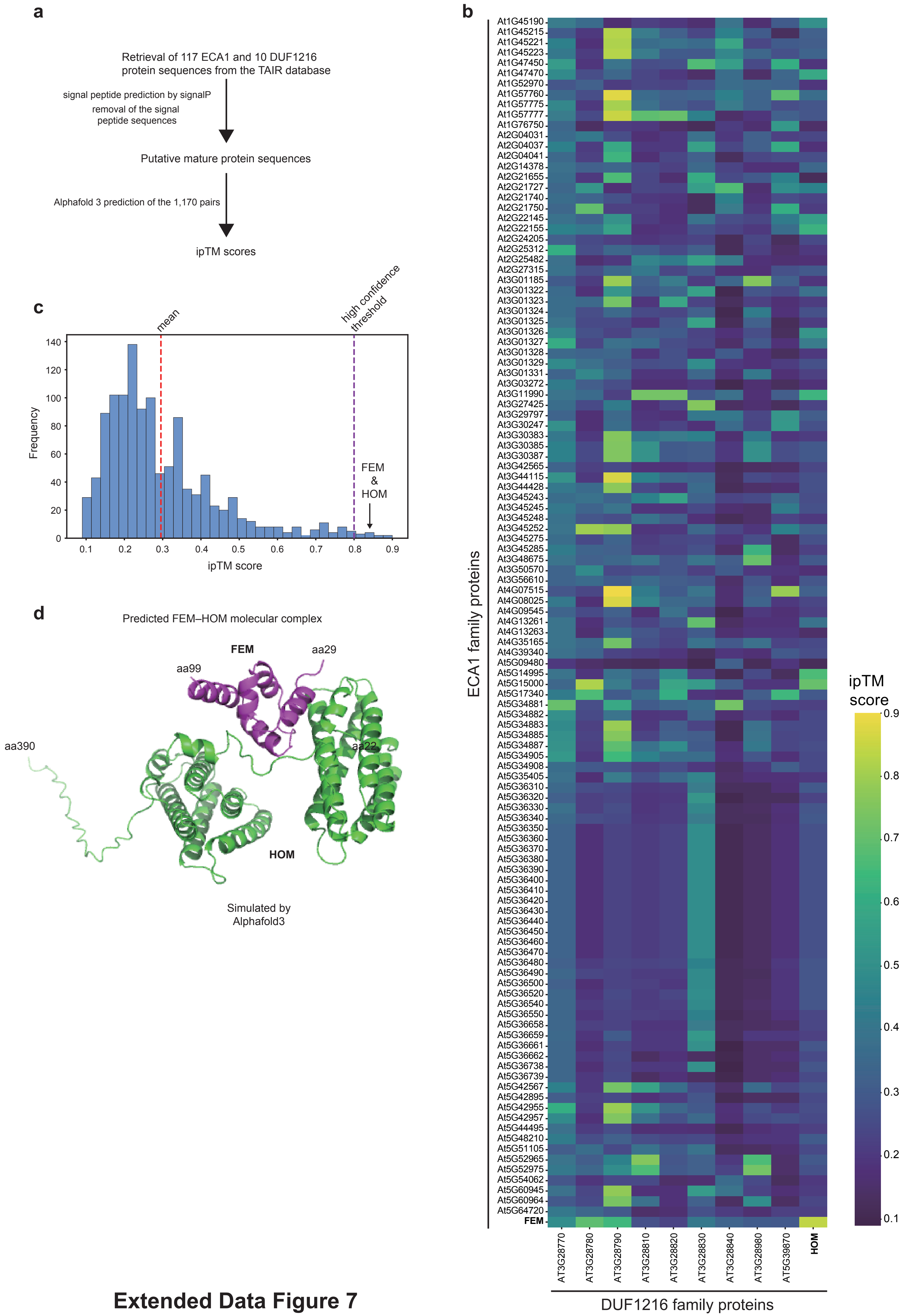


### Extended Data Figure 7

**a** World map

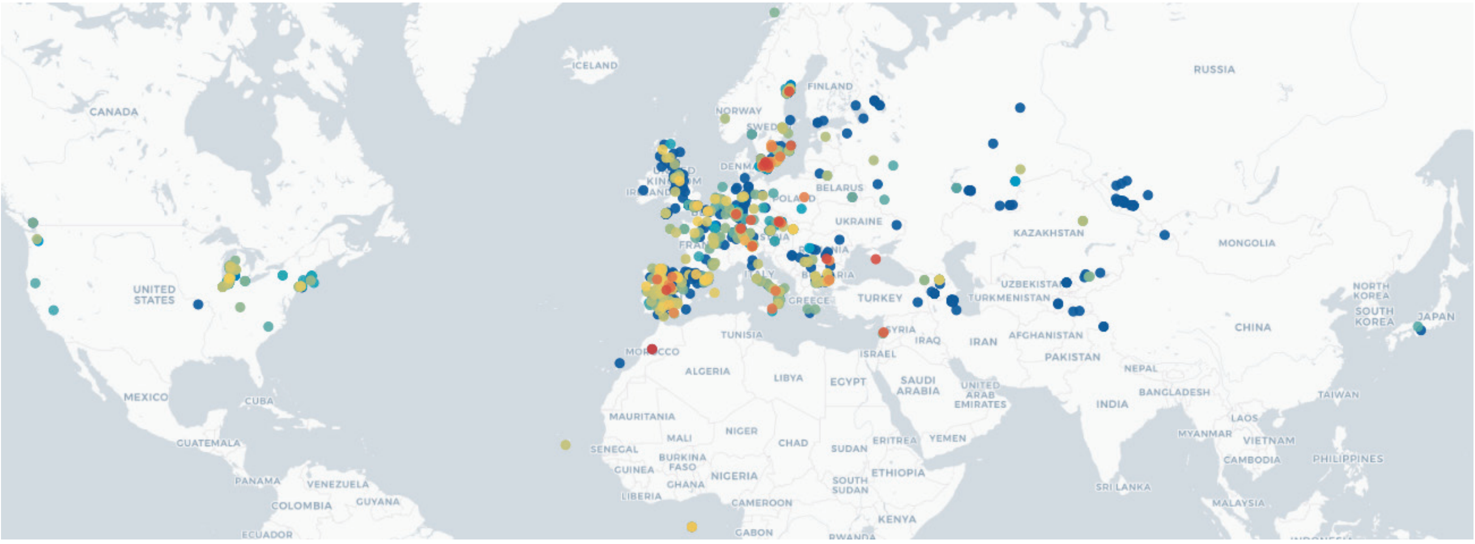


**b** Europe-Africa

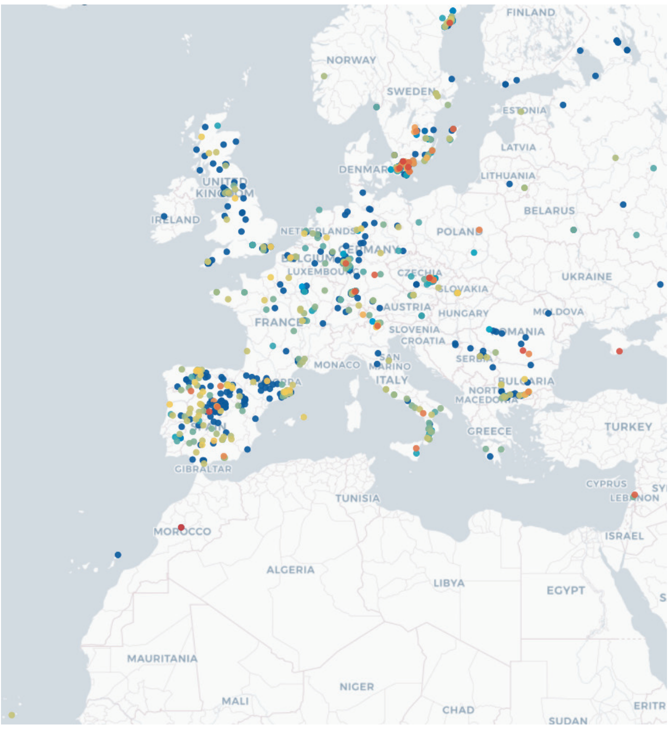


**c** Central Europe

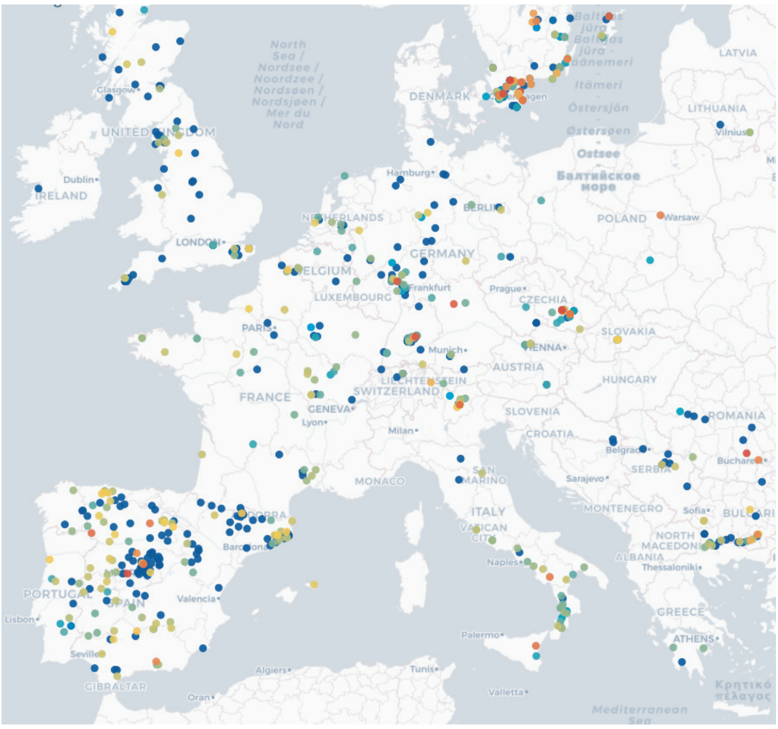


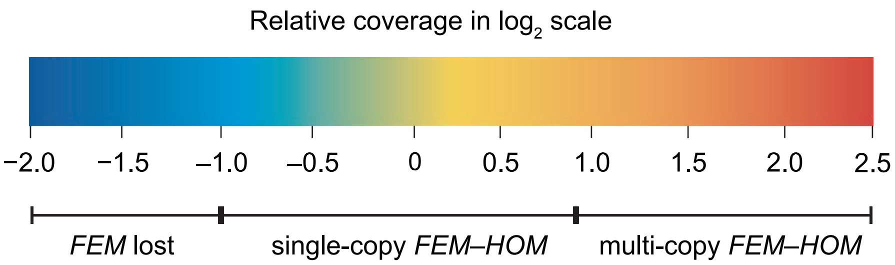


### Extended Data Figure 8

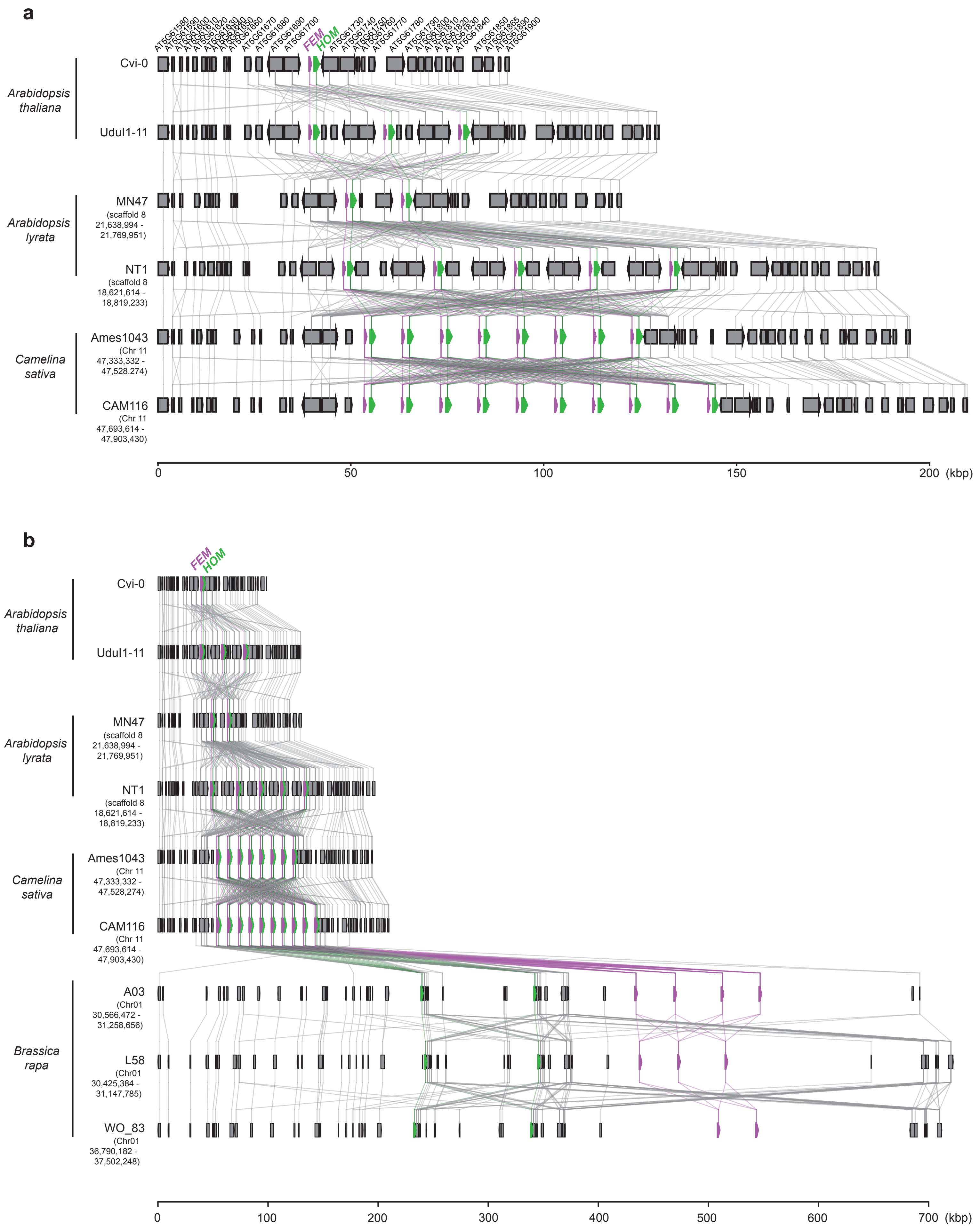


### Extended Data Figure 9
